# Prior expectation enhances sensorimotor behavior by modulating population tuning and subspace activity in the sensory cortex

**DOI:** 10.1101/2022.12.04.516847

**Authors:** JeongJun Park, Seolmin Kim, HyungGoo R. Kim, Joonyeol Lee

**Author notes:** These authors contributed equally to this work (JeongJun Park, Seolmin Kim). Proofs and correspondence to: Joonyeol Lee, Department of Biomedical Engineering, Sungkyunkwan University, 2066 Seobu-ro, Jangan-gu, Suwon-si, Gyeonggi-do, 16419, Republic of Korea. Author contributions: Conceptualization, JP, SK, and JL; Methodology, JP, SK, HRK, and JL; Investigation, JP and SK; Formal Analysis, JP, SK, and JL; Writing – Original Draft, JP, and JL; Writing – Review and Editing, JP, SK, HRK, and JL; Funding Acquisition, JL. ***Data Availability:*** The data and the code supporting this study’s findings can be accessed from the link, https://semoconlab.com/codes/bayesian-area-mt/.

## Abstract

Prior knowledge facilitates our perception and goal-directed behaviors in the dynamic world, particularly when sensory input is lacking or noisy. However, the neural mechanisms underlying the improvement in sensorimotor behaviors by prior expectations remain unknown. In this study, we examine the neural activity in the middle temporal (MT) area of visual cortex while monkeys perform a smooth pursuit eye movement task with prior expectation of the visual target’s motion direction. Prior expectations discriminately reduce the MT neural responses depending on their preferred directions, only when the sensory evidence is weak. This response reduction effectively sharpens neural population direction tuning. Simulations with a realistic MT population demonstrate that sharpening the tuning explains both the biases and variabilities in smooth pursuit, thus suggesting that neural computations in the sensory area alone can underpin the integration of prior knowledge and sensory evidence. State-space analysis further supports this by revealing neural signals of prior expectation in the MT population activity that correlate with behavioral changes.

## Introduction

When interacting with our environment, we quickly adapt to its dynamic changes and adjust our actions based on the quality of sensory information. If the available sensory information is sufficiently precise to guide behaviors, our actions can depend solely on sensory input. However, if the sensory information is imprecise, we combine it with prior knowledge to improve our behavioral responses. In particular, we use the reliability of each piece of information as a weight when combining multiple pieces of information. This precision-weighted information integration explains the fundamental approaches to our interactions with the environment.

The Bayesian inference framework has successfully explained information integration in perceptual decision-making ^1–4^, multisensory integration ^5–9^, and sensorimotor behaviors ^10–13^. The neural implementation of Bayesian inference has also been demonstrated in studies on sensory and motor functions ^5,14–22^. Notably, preparatory activity in the smooth eye movement region of the frontal eye field (FEF_SEM_) represented the prior distribution for incoming motion speed, whereas evoked activity reflected a Bayesian estimate of the stimulus speed during smooth pursuit eye movements ^14,23,24^. However, the effect of prior expectations on neural sensory representation has achieved insufficient consensus ^25^. Human neuroimaging studies have suggested that prior expectations reduce neural activity ^26–31^ and modulate sensory representation in the early visual cortex ^31–33^. By contrast, previous single-cell electrophysiological data have shown that prior expectations have no effect on sensory motion representation in the middle temporal (MT) area of visual cortex ^34^. Therefore, although some experimental evidence supports the neural origins of Bayesian inference, the effects of prior expectations on cortical sensory neurons are scarce and contradictory.

In this study, single neuronal activity was recorded from area MT of two rhesus monkeys performing a smooth pursuit eye movement task wherein the strength of the sensory motion information and prior knowledge of motion direction were controlled. Consistent with the findings of our previous behavioral study ^35^, the variation in pursuit directions across trials was reduced by prior expectations only when the sensory input was weak and imprecise. The neural recordings indicated that prior expectations systematically reduced the MT neural responses in a manner that sharpened the population direction tuning curve only when the sensory evidence was weak. The simulation of a realistic MT population activity demonstrated that this systematic reduction of neural responses accounted for the reduction in behavioral variability observed in the initiation of smooth pursuit eye movement. The integration of prior knowledge with sensory information in area MT and its role in behavioral enhancement were additionally supported by the targeted dimensionality reduction analysis of neural population activity. The neural state of the MT population represented prior information and its temporal evolution accounted for the dynamic modulation of behavioral variability reduction observed in smooth pursuit initiation.

## Results

In this study, we aimed to identify the neural correlates of prior expectations in area MT using a behavioral paradigm that allowed the direct manipulation of prior information with an identical sensory stimulus ^35^. We measured the spiking activity of single neurons in area MT of two rhesus monkeys while performing a smooth pursuit eye movement task (Fig. 1a). Each trial began with a fixation spot appearing at the center of the monitor screen. A random-dot kinematogram appeared after the fixation duration, and all dots of the stimulus moved in a direction within a fixed window for 100 ms (local motion). Following the local motion, the dot patch smoothly moved in the same direction as the local motion, and the monkeys had to maintain their gaze on the visual target during its movement. A liquid droplet was delivered as a reward for the correct behavior at the end of the trial. The expectation of motion direction was manipulated in two blocks: “wide prior block” and “narrow prior block” (Fig. 1b). Beginning with a randomly selected block, the two prior blocks were alternated. In the wide prior block, the direction of the pursuit target was randomly selected from three widely distributed directions, which were 120° apart from each other. The number of trials in all three directions was the same. In the narrow prior block, the target direction was one of the three narrowly distributed directions, wherein the difference between the central direction and the others was 15°. The central direction was presented twice as often as the other directions. Therefore, the monkeys had prior expectations of the central direction only in the narrow prior block. Notably, the central direction of the narrow prior block, termed “prior direction,” was also included in the wide prior block to enable comparison of the eye movements and neural responses between the two blocks under the identical stimulus condition. The prior direction was set by considering the preferred directions of the recorded MT neurons. To change the sensory evidence, two different stimulus types, “high contrast” and “low contrast”, were randomly interleaved; these had different luminance contrasts (100 % and 12 % for Monkey A; 100 % and 8 % for Monkey B) and coherence (without and with random-walk noise) of dots in both blocks (Fig. 1c, see Methods). Additionally, in each prior block, the smooth pursuit trials were randomly interrupted using direction tuning trials to measure the direction tuning curves of the neurons being recorded (see Methods for details). Through direction tuning measurements, neural responses to 12 motion directions (from 0° to 330° in 30° increments) were obtained and the direction tuning functions of individual neurons were estimated using a Gaussian or circular Gaussian function.

**Fig. 1:**
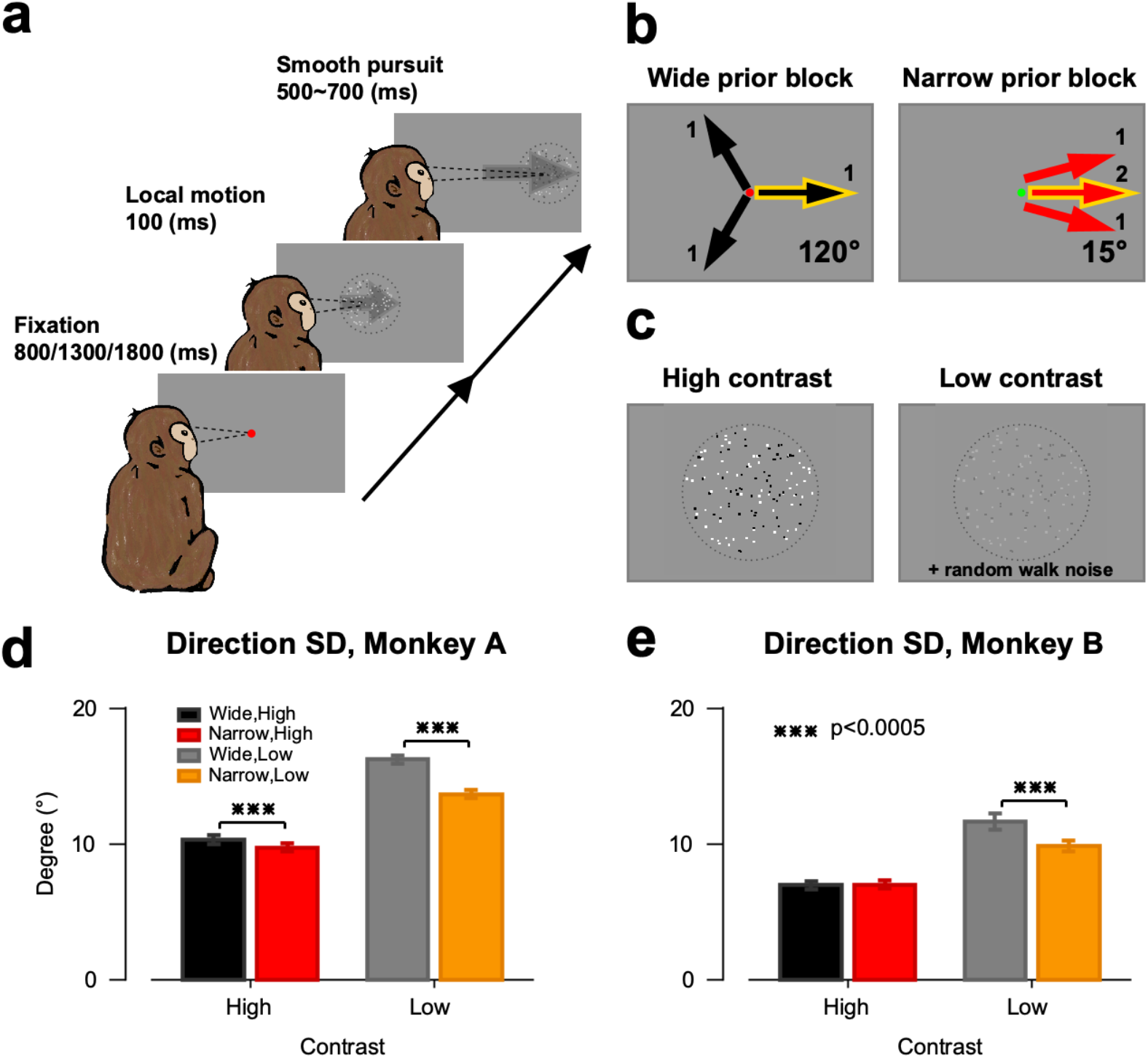
Experimental design and task performance. **a** Trial timeline; Each trial begins with the monkeys fixating their eyes on a spot at the center of the monitor screen. After a randomized fixation duration (800, 1300, or 1800 ms), a random-dot kinematogram appears at the center of the screen or 1°–2° displaced from the center to the opposite direction of the target direction, and all the dots inside the fixed invisible window move in the target direction for 100 ms. When the fixation spot disappears after the local motion, the dot patch smoothly moves in the same direction as the local motion for 500–700 ms, and the animals have to maintain their gaze at the center of the moving patch. **b** Block design; The monkeys’ expectations of the direction of the pursuit target are manipulated by presenting different sets of three directions in different proportions in two types of blocks. Both blocks have a common direction (“prior direction”), and the angle between that direction and the other directions is 120° and 15° in the wide and narrow prior block, respectively. The number of trials for the prior direction is the same as for the other directions in the wide prior block but twice that in the narrow prior block. The wide and narrow prior blocks are distinguished by red and green fixation points, respectively. **c** Stimulus type; To control for sensory evidence, two different types of random-dot patches are randomly interleaved as visual stimuli in both blocks. The high contrast stimuli has 100 % luminance contrast and dot coherence, whereas the low contrast stimuli has 12 % (Monkey A) or 8 % (Monkey B) luminance contrast and random-walk noise in dot motion. **d, e** Mean standard deviation (SD) of pursuit directions for the prior direction target. The black and red bars show the mean SD for the high-contrast cases in the wide and narrow prior blocks, respectively; the gray and yellow bars show the mean SD for the low contrast cases in the wide and narrow prior blocks, respectively. The error bars indicate the standard error of the mean.

### Prior expectation reduces the variability of pursuit eye movements

Our previous study ^35^ demonstrated that prior expectation of motion direction reduces the variation in pursuit directions for moving stimuli. We reconfirmed the effect of the direction prior to the smooth pursuit eye movements. To compare the variability in pursuit eye movements in the same direction between the wide and narrow prior blocks, the standard deviation (SD) of pursuit directions for prior direction was calculated in each block during the open-loop period (100 ms from pursuit onset ^36^) to exclude any impact of feedback signals (see Methods). Figs. 1d, e show the mean SD of pursuit directions from the two monkeys (59 days’ data for Monkey A and 67 days’ data for Monkey B). In Monkey A, the mean SD of the narrow prior block was smaller than that of the wide prior block in both high and low contrast cases (mean SD in the wide and narrow prior blocks was 10.28° vs. 9.72° for high contrast stimuli, paired t-test, t(58) = 3.72, *p* = 4.46 × 10^−4^, 16.20° vs. 13.64° for low contrast stimuli, paired t-test, t(58) = 12.70, *p* = 2.18 × 10^−18^). By contrast, the difference was significantly larger when the stimulus contrast was low (mean SD difference between two prior blocks in high and low contrast cases: 0.56 vs. 2.57, paired t-test, t(58) = –8.39, *p* = 1.35×10^−11^). In Monkey B, the difference in SD between the two blocks was significant only for low contrast stimuli (mean SD in the wide and narrow prior block: 6.94° vs. 7.00° for the high contrast stimuli, paired t-test, t(66) = –0.61, *p* = 0.54, 11.66° vs. 9.83° for low contrast stimuli, paired t-test, t(66) = 6.94, *p* = 2.33×10^−9^). These results demonstrate that prior expectation of motion direction decreases the variation in pursuit directions, particularly when the sensory motion of the pursuit target is weak and imprecise, which is consistent with the prediction of Bayesian inference ^35^.

### Prior expectation systematically reduces MT neural responses only when sensory input is weak and imprecise

Recent human functional magnetic resonance imaging studies have reported that prior expectation of a stimulus feature reduces neural responses in the primary visual cortex ^29,31^ and enhances neural representations in early visual areas ^31,32^. A more recent magnetoencephalography study supported the role of expectation in perceptual processes by demonstrating that expectations modulate the neural representation in early sensory processing ^33^. In this study, we tested whether the responses of the MT neurons were modulated by prior expectations. To determine the neuronal correlates underlying the effect of prior expectation on the variation in eye movements, we recorded 257 well-isolated single neuronal activities (137 for Monkey A and 120 for Monkey B) from area MT while the monkeys performed the task. The peristimulus time histograms (PSTHs) in Figs. 2a, b show the firing rate of MT neurons in the prior direction as a function of time. When the stimulus contrast was low, the firing rate in the narrow prior block was lower than that in the wide prior block in the initial neural responses of both monkeys (cluster-based permutation test, alpha = 0.05, Monkey A: 61–129 ms; Monkey B: 101–199 ms from the pursuit target onset). However, no common decrease in the firing rate was observed between the two monkeys when the stimulus contrast was high. We particularly focused on early neural responses because it was difficult to determine whether the neural response reduction occurring after the open-loop period (up to 100 ms after pursuit onset or 200 ms after motion onset) was induced by prior expectation or the feedback signal for the difference between the target and eye movements. To further quantify the reduction in neural responses, we calculated the firing rates during the initial 100 ms from the spike latency of each neuron in response to the pursuit target. The insets in Figs. 2a, b depict the mean firing rates of the MT neurons in the time window under the two prior conditions. As shown in the PSTHs, the early MT neuronal responses after the spike latency were smaller in the narrow prior block than in the wide prior block only when the stimulus contrast was low (mean firing rate of the wide and narrow prior block in Monkey A was 43.37 vs. 42.64 for the high contrast case, paired t-test, t(135) = 1.82, *p* = 0.07; 25.21 vs. 23.17 for the low contrast case, paired t-test, t(110) = 4.53, *p* = 1.51×10^−5^, that in Monkey B: 49.86 vs. 49.38 for the high contrast case, paired t-test, t(116) = 1.15, *p* = 0.25; 25.45 vs. 23.98 for the low contrast case, paired t-test, t(109) = 3.35, *p* = 0.001). These results indicate that prior expectation of motion direction reduces the responses of MT neurons only when sensory motion is weak and imprecise.

**Fig. 2:**
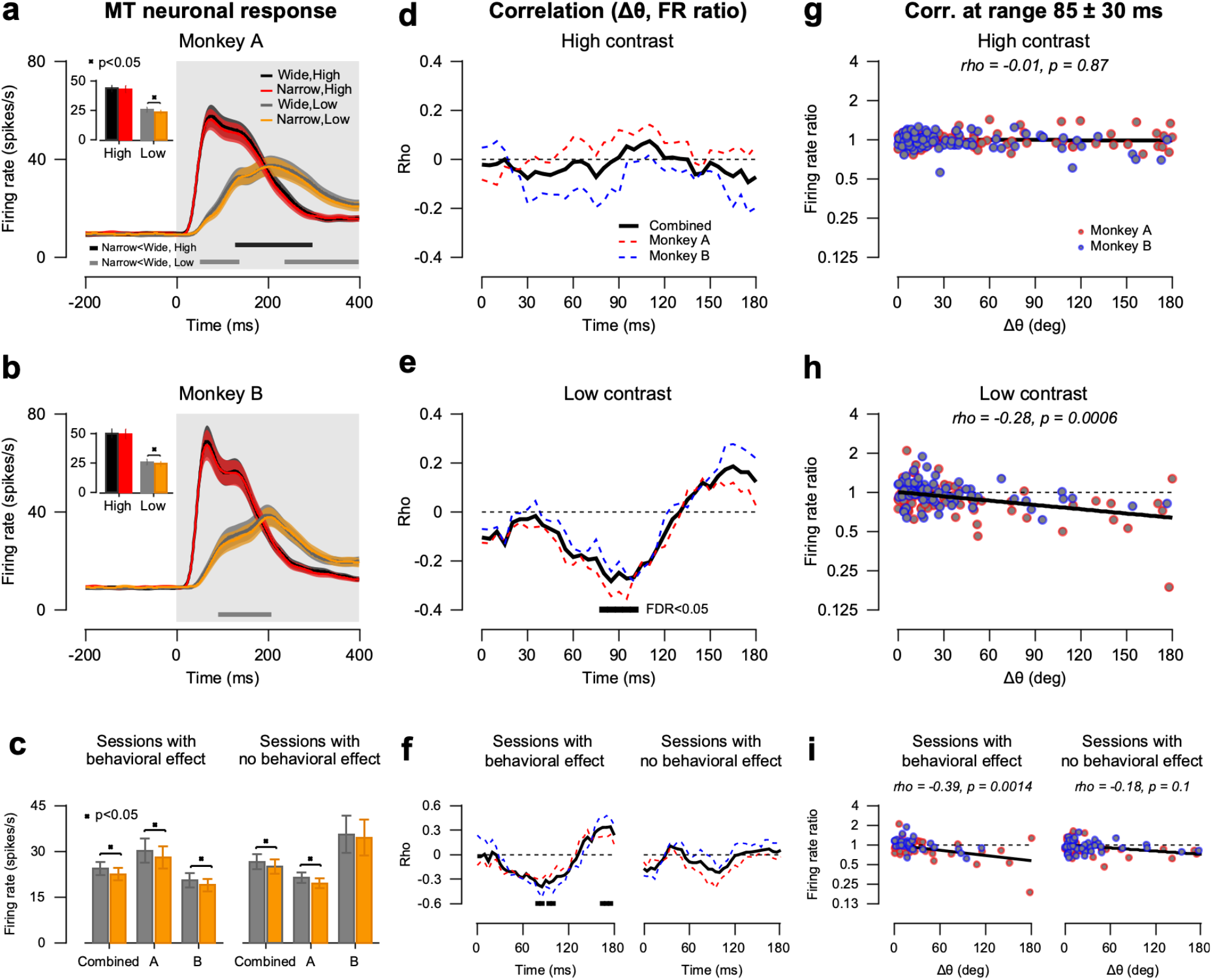
Effect of prior expectation on the firing rate of MT neurons. **a–c** Each colored peristimulus time histograms (PSTHs) shows the firing rate of the MT neurons for the prior direction as a function of time relative to the stimulus onset in each condition (black: wide prior, high contrast; red: narrow prior, high contrast; gray: wide prior, low contrast; yellow: narrow prior, low contrast). The bar graphs show the mean firing rates during the initial 100 ms time window from spike latency. **d–f** Correlation between the difference in the prior and preferred directions (Δθ) and the firing rate ratio in the narrow prior block to that in the wide prior block (firing rate ratio). The lines show the correlation coefficient over time relative to the stimulus onset with a ±30 ms window and 5 ms timestep; the red, blue, and black lines are from Monkey A, Monkey B, and the combined dataset of Monkeys A and B, respectively. **g–i** Correlation between Δθ and firing rate ratio in the time window of 55-115 ms from stimulus onset in the combined dataset. Each point on the plot represents one MT neuron in the combined dataset. The black solid lines show the linear regression relationship between Δθ and the firing rate ratio. **c, f, i** The MT neuronal responses for low contrast stimuli on the days where the SD in pursuit directions in narrow prior block are significantly smaller than those in wide prior block, and the responses on the other days with no difference in SD between the two prior blocks.

The small and uniform decrease in neural responses may not account for the decrease in behavioral variability. However, if the response reduction depends on the difference between the preferred directions of the neurons and the prior direction, it can enhance the neural representation of the motion direction by modulating the population direction tuning function. The reduction in MT neuronal responses by prior expectation indeed varied according to the relationship between the prior direction and the preferred direction of each neuron. We calculated the ratio of the firing rate in the narrow prior block to that in the wide prior block to account for the different firing rates between the two prior blocks. We then estimated the correlation coefficient between the firing rate ratio (FR ratio) and the difference in the prior and preferred directions (Δθ). In this analysis, neurons whose direction tuning functions have been relatively well defined (fitted tuning function explained more than 50% of data variance) were included for the robust estimation of the preferred direction of each neuron. With these additional criteria, 181 (n = 95 for Monkey A and 86 for Monkey B) and 144 (n = 75 for Monkey A and 69 for Monkey B) neurons were retained for the high and low contrast cases, respectively. The correlation from each monkey and that from the combined dataset are plotted as a function of time relative to the pursuit target onset (Figs. 2d, e). The correlation did not differ from zero when the stimulus contrast was high (Fig. 2d). However, when the stimulus contrast was low, the correlation decreased after the stimulus appeared, and it was significantly lower than zero between 80 and 100 ms from the stimulus onset in the combined dataset (Fig. 2e, time windows for calculation: ± 30 ms at each time point of 0–180 ms in 5 ms intervals, Spearman correlation, false discovery rate (FDR) corrected, alpha = 0.05). Figs. 2g, h show the relationship between the FR ratio and Δθ at 85 ± 30 ms when they were the most negatively correlated in the low contrast stimuli case (Spearman correlation, *ρ* = –0.01, *p* = 0.87 for high contrast; *ρ* = –0.284, p = 0.00059 for low contrast). This means that as the preferred direction of an MT neuron was farther from the prior direction, the neuron’s response to the prior direction decreased further when the sensory input was weak. This relationship was still significant for neurons whose preferred direction was less than 90° apart from the prior direction, although the overall correlation was smaller (*ρ* = –0.18, *p* = 0.0477 in low contrast condition).

The animals’ behavioral performace on the pursuit task varied from day to day. The difference in pursuit direction SD between the wide and narrow prior blocks might depend on changes in the use of prior expectation. In both cases, the difference in pursuit directional variation between the two blocks indicates whether the behavioral effect of prior expectation was strong or weak that day. To further examine whether neural modulation is tightly linked with smooth pursuit behavior, we divided the experimental sessions into two groups based on whether the variability of pursuit direction was significantly reduced by prior expectations. We compared the reduction in neural activity between the two groups, sessions with vs. without behavioral effect by prior expectation (number of neurons for significant vs. nonsignificant direction SD reduction in the low contrast stimuli case: n = 62 for 29 days vs. n = 75 for 30 days in Monkey A; n = 83 for 35 days vs. n = 37 for 32 days in Monkey B; two-sample F-test, alpha = 0.05). When the pursuit direction variance was significantly reduced by prior expectation, the responses of MT neurons also decreased, and the response reduction depended on the difference between the prior and preferred directions, as we observed in the entire dataset. However, when the reduction in pursuit direction variance was insignificant, even if the prior expectation reduced MT responses, the reduction was not correlated with Δθ (Fig. 2c: mean firing rate in wide vs. narrow prior block of Monkey A: 30.13 vs. 27.93 on the days with significant behavioral effect, paired t-test, t(61) = 2.73, *p* = 0.009; 21.32 vs. 19.41 on the days with non-significant behavioral effect, paired t-test, t(74) = 3.83, *p* = 3.04×10^−4^; that of Monkey B: 20.52 vs. 18.88 on the days with significant behavioral effect, paired t-test, t(82) = 2.99, *p* = 0.004; 35.56 vs. 34.48 on the days with nonsignificant behavioral effect, paired t-test, t(36) = 1.51, *p* = 0.14, Figs. 2f, i: Spearman correlation between FR ratio and Δθ in 85 ± 30 ms from the stimulus onset: *ρ* = –0.39, *p* = 0.0014 on the days with significant behavioral effect; *ρ* = –0.18, p = 0.1 on the days with nonsignificant behavioral effect). This relationship indicates that the systematic reduction of neural responses was controlled by the strength of the prior expectation that monkeys employed, thus suggesting that the decrease in pursuit direction variability by prior expectation may be driven by the systematic response reduction of cortical sensory neurons.

### Decoded direction information from the simulated MT responses can account for the behavioral modulation by prior expectation

The systematic reduction in neuronal responses, which depends on the difference between the prior direction and the preferred directions of MT neurons, can change the shape of the population direction tuning curve in area MT. If the prior expectation is congruent with stimulus motion, it can sharpen the population direction tuning curve and enhance the neural representation of motion direction (Fig. 3a). To test whether the observed response reduction has a noticeable impact on the quality of sensory neural representation and the resulting behavioral performance, neuronal activity was simulated using realistic properties obtained from our MT recordings. Using the estimated parameters of the circular Gaussian functions for direction tunings of recorded MT neurons, we simulated 3600 neuron responses whose preferred directions were randomly selected from a uniform distribution bounded by –180° and 180° (see Methods for details). The measured tuning shape, average firing rate, Fano factor, and the inter-neuronal correlation were considered in the simulation; therefore, the simulated neurons had realistic neural response properties (see Methods and Supplementary Fig. 1 for further details). Subsequently, the effect of prior expectation on neural activity was modeled by applying slopes of a linear regression between Δθ (preferred–prior direction) and the log firing rate ratio (narrow/wide prior) that was estimated from the recordings (regression coefficient at 85 ± 30 ms from the pursuit target onset: –1.06×10^−4^ for the high contrast case, *p* = 0.60; –0.0025 for the low contrast case, *p* = 1.01×10^−6^). The simulated population responses for the high contrast stimuli did not differ between the two prior blocks because the slope of regression was close to zero. However, the simulated population tuning curves for the low contrast stimuli became narrower in the narrow prior condition because of the negative relationship between Δθ and the firing rate ratio. Figs. 3b, c show examples of simulated responses for high and low contrast cases, respectively, when the target and prior directions are 0°.

**Fig. 3:**
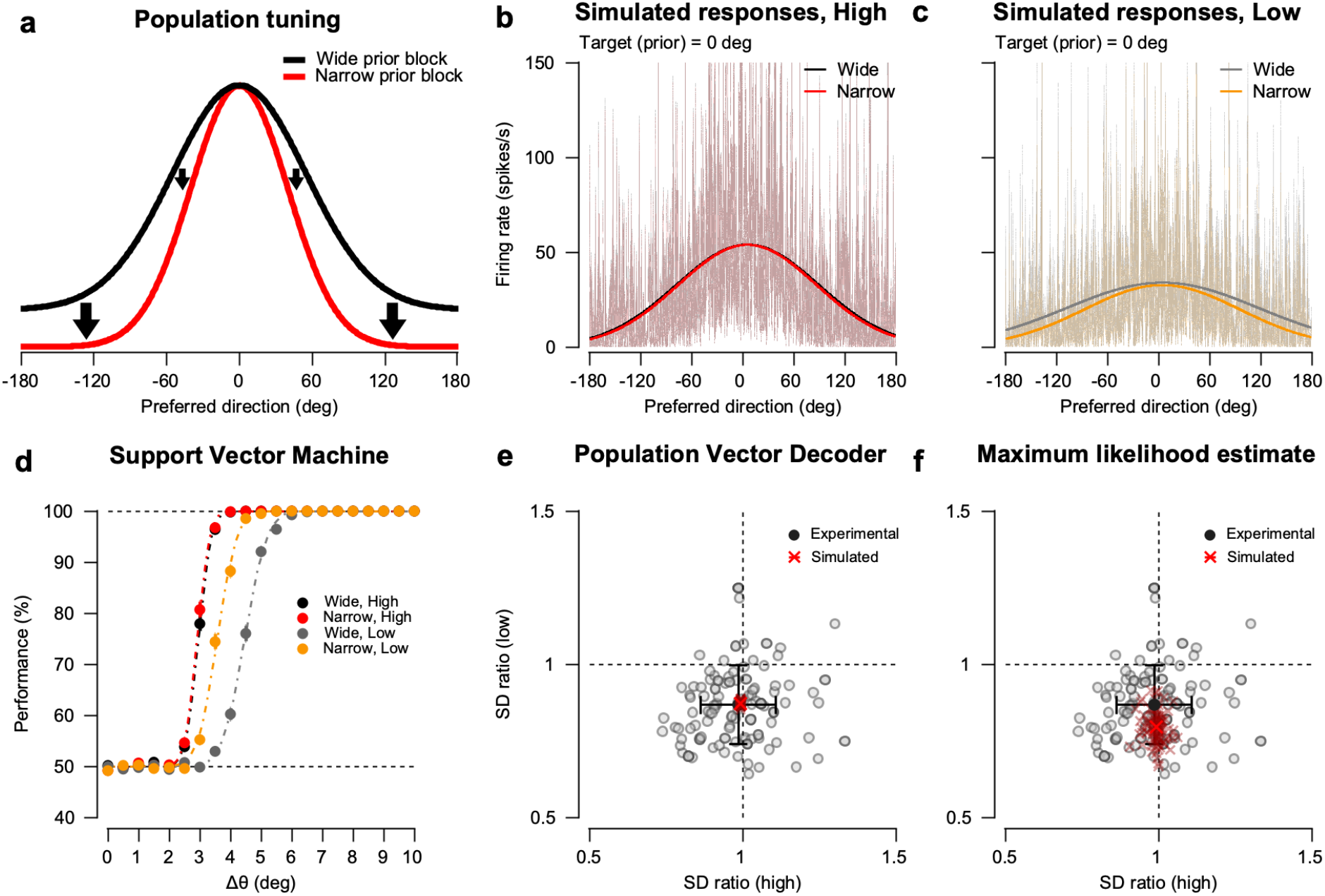
In silico simulations to decode population direction information. **a** Schematic of the change in the population tuning curve in area MT by prior expectation. The black and red curves exhibit the population direction tuning of the model MT neurons in the wide and narrow prior blocks when the target direction and prior direction are 0°; responses to the target as a function of neuron’s preferred direction. Based on the experimental results, when the sensory evidence is weak, prior expectations reduce the MT neuronal responses proportionally to the difference between the prior direction and each neuron’s preferred direction. This sharpens the shape of the population tuning curve. **b, c** Example of simulated MT population responses under high/low contrast and wide/narrow prior. Each point on the thin line plots represents the firing rate of each model neuron in a single trial with the target (and the prior) direction of 0°. Thick curves show the mean population responses across the trials. **d** Performance of support vector machine (SVM) in discriminating between two directions as a function of direction difference (Δθ). The direction discrimination performances of the SVM classifier (colored circles) at every Δθ were fitted to a cumulative Gaussian function (dot-dashed line). **e, f** Estimation of direction SD ratio between the narrow prior block and the wide prior block in the experimental and simulated datasets. Gray circles show the SD ratio from individual smooth pursuit direction variability in each session of the two monkeys, and black circles represent the mean of the SD ratios. The light red crosses show the SD ratio estimated from the population responses in each simulation using a population vector decoder (**f**, left) or using a maximum likelihood estimation method (**f**, right), and the thick red crosses represent the mean SD ratio.

Using these simulated MT neural responses, we tested whether the neural direction discrimination could be improved by the prior factor. Neural population responses to two different motion directions were simulated and a binary linear support vector machine (SVM) discriminated the two directions, which were 0°, and a direction from 0° to 10° with step sizes of 0.5° (see Methods for details). In this simulation, the target and prior directions were identical, and the prior factor was applied to each direction separately such that the population tuning curve became centered on each stimulus direction (Figs. 3b, c). Therefore, the prior factor only influenced the level of noise in the neural direction representation. When the difference between the two directions was small, the SVM classifier could not discriminate between the directions (Fig. 3d, direction difference <2°). However, its performance rapidly increased as the direction difference increased. The performance of the SVM classifier was better in the high contrast condition than in the low contrast condition (mean of the cumulative Gaussian function in high and low contrast conditions of wide prior block : 2.97 vs. 4.45, paired t-test, t(99) = –42.26, *p* = 3.77×10^−65^; that of narrow prior block: 2.92 vs. 3.57, paired t-test, t(99) = –19.79, *p* = 3.52×10^−36^), where the simulated population responses were more robust. However, the effect of the prior factor in the high contrast condition was negligible (mean of the cumulative Gaussian function for high contrast stimuli: 2.97 (wide) vs. 2.92 (narrow), paired t-test, t(99) = 1.66, *p* = 0.1). By contrast, the effect of the prior factor on neural direction discrimination was significant in the low contrast condition. When the classifier was trained and tested on the simulated neural responses in the low contrast, narrow prior condition, the neural direction discrimination was significantly better than that when the classifier was trained in the wide prior condition (mean of the cumulative Gaussian function for low contrast stimuli: 4.45 (wide) vs. 3.57 (narrow), paired t-test, t(99) = 26.04, *p* = 4.28×10^−46^). These findings suggest that the systematic response reduction observed in MT neuronal activity to low contrast stimuli has a significant effect on enhancing the neural motion direction representation, probably by reducing population-level neural noise.

The SVM demonstrated that prior expectations improve the signal-to-noise ratio in the population-level neural representation of the motion direction. To determine the behavioral enhancement by prior expectation can be accounted for by the modulation of population direction tuning in area MT, we compared the neuronal variability (i.e., variance in neural direction representation) with behavioral variability (i.e., variance in pursuit direction). For this comparison, a population vector decoder (PVD) ^37^ and maximum likelihood estimate (MLE) ^38–40^ were used to directly measure the trial-by-trial variability of neural direction estimates (see Methods for details). The direction in each trial was decoded from the simulated population response in a 0° motion direction, and the SD of the direction estimates was calculated across trials. To estimate the extent to which the direction variability was reduced by prior expectation, we computed the ratio of the SDs in the narrow and wide prior blocks (SD_narrow prior_/SD_wide prior_). The mean SD ratio of the direction estimates from simulations was smaller than 1, which suggests that the simulated SD in the narrow prior block was smaller than that in the wide prior block and the effect was stronger when the stimulus contrast was low (Figs. 3e, f, mean SD for high contrast stimuli: 0.994 and 0.993, one-sample t-test, t(99) = 2.74×10^4^ and 384.16, *p* = 0 and 6.39×10^−159^; mean SD for low contrast stimuli: 0.87 and 0.80, one-sample t-test, t(99) = 1.37×10^3^ and 157.58, *p* = 1.70×10^−213^ and 1.12×10^−120^ from PVD and MLE, respectively). This was consistent with the findings from the behavioral data (Figs. 3e, f, mean SD for high contrast stimuli: 0.99, one-sample t-test, t(99) = 89.92, *p* = 1.83×10^−115^; mean SD for low contrast stimuli: 0.87, one-sample t-test, t(99) = 75.47, *p* = 4.00×10^−106^). Notably, the SD ratio of the directions estimated by PVD was almost the same as the experimental SD ratio (experimental vs. simulated SD ratio in the high contrast case: 0.9863 vs. 0.9936, pooled t-test, t = –0.60, *p* = 0.55; those in low contrast case: 0.8677 vs. 0.8721, pooled t-test, t = –0.34, *p* = 0.73). This close correspondence between the experimental and simulated results suggests that the modulation of the shape of the population direction tuning curve can fully account for the reduction in behavioral variation in smooth pursuit initiation.

As demonstrated before ^35^, prior expectations introduced an increase in behavioral bias as well as a decrease in behavioral variability during the pursuit task, only when the sensory input was weak and imprecise. The directions of smooth pursuit eye movements to the outer target directions were skewed toward the prior direction; the ratio between the angular difference of two outer target directions and that of the corresponding pursuit directions was smaller than 1 in the low contrast cases (Supplementary Figs. 2a, b, 0.997 and 0.801 for high and low contrast stimuli, paired t-test, t(42) = –0.25 and –9.66, *p* = 0.81 and 1.82×10^−15^). To test if the bias can also be explained by the expectation-induced modulations in the population tuning, we simulated population responses in ±15° motion direction when the prior direction was 0° and decoded using PVD and MLE, respectively (see Supplementary Fig. 2 and Methods for details). Consistent with the experimental results, the simulation showed that the estimated directions were biased toward the prior direction, and this effect was significantly stronger when the stimulus contrast was low (Supplementary Figs. 2c–f, the ratio between the two differences for high contrast stimuli: 0.994 and 0.987, one-sample t-test, t(99) = –244.59 and –57.47, *p* = 1.57×10^−139^ and 7.54×10^−78^; that for low contrast stimuli: 0.84 and 0.75, one-sample t-test, t(99) = –261.12 and –1.12×10^3^, *p* = 2.45×10^−142^ and 8.29×10^−205^ from PVD and MLE, respectively). These results demonstrate that the systematic change in the population response of MT neurons by prior expectations can explain the bias and variability reduction in pursuit directions. Additionally, this further suggests that modulation of MT neural responses is sufficient to explain Bayesian inference in smooth pursuit eye movements.

### Systematic reduction in neuronal responses occurs only in an active behaving condition

As suggested earlier, prior expectations improve pursuit behavior by sharpening the shape of the population neural direction tuning. However, the systematic response reduction might result from passive response modulation by the repetitive presentation of similar stimuli. That is, these observations could originate from neural adaptation ^41^, and not prior expectation (although they are tightly correlated). To address this possibility, we measured the direction tuning responses of MT neurons in the middle of each prior block. Even if the same motion stimuli were presented, the monkeys did not have to use prior knowledge because the animals passively fixed their eye gazes on the central fixation point. By contrast, changes in the passive properties of individual neurons that were induced by the different statistical distributions of motion stimuli across blocks could be revealed. To measure direction tuning, we used identical visual stimuli as the pursuit targets and placed them in the center of the receptive fields of the neurons. Directions were randomly selected from 12 directions (0°, 30°, …, 300°, 330°) in each presentation. To obtain the mean direction tuning curve across all neurons from the two monkeys, we pooled the direction tuning responses by aligning them relative to the preferred direction of each neuron. Subsequently, we fit the realigned mean neural responses to the Gaussian function and compared the averaged direction tuning function between the wide and narrow prior blocks. As a result, there was no significant difference in mean direction tuning between the two blocks (Figs. 4a–c, the height of the Gaussian function in the wide and narrow prior blocks: 51.08 and 50.48 for high contrast stimuli, 33.49 and 33.29 for low contrast stimuli in the combined dataset; 45.05 and 44.54 for high contrast stimuli, 29.29 and 29.06 for low contrast stimuli in Monkey A; 58.26 and 57.63 for high contrast stimuli, 38.72 and 38.56 for low contrast stimuli in Monkey B). Additionally, the tuning function of individual neurons was estimated by fitting Gaussian or circular Gaussian functions (selected better function between the two for each neuron). The amplitudes and half-widths of the tuning curves were not different across the blocks in both high and low contrast cases (paired t-test, mean tuning amplitude: 55.48 vs. 55.05 for high contrast, t(164) = 0.86, *p* = 0.39, 32.03 vs. 31.88 for low contrast, t(164) = 0.32, *p* = 0.75, mean tuning half-width: 85.11 vs. 86.71 for high contrast, t(164) = –0.82, *p* = 0.41, 87.88 vs. 87.08 for low contrast, t(164) = 0.18, *p* = 0.86).

**Fig. 4:**
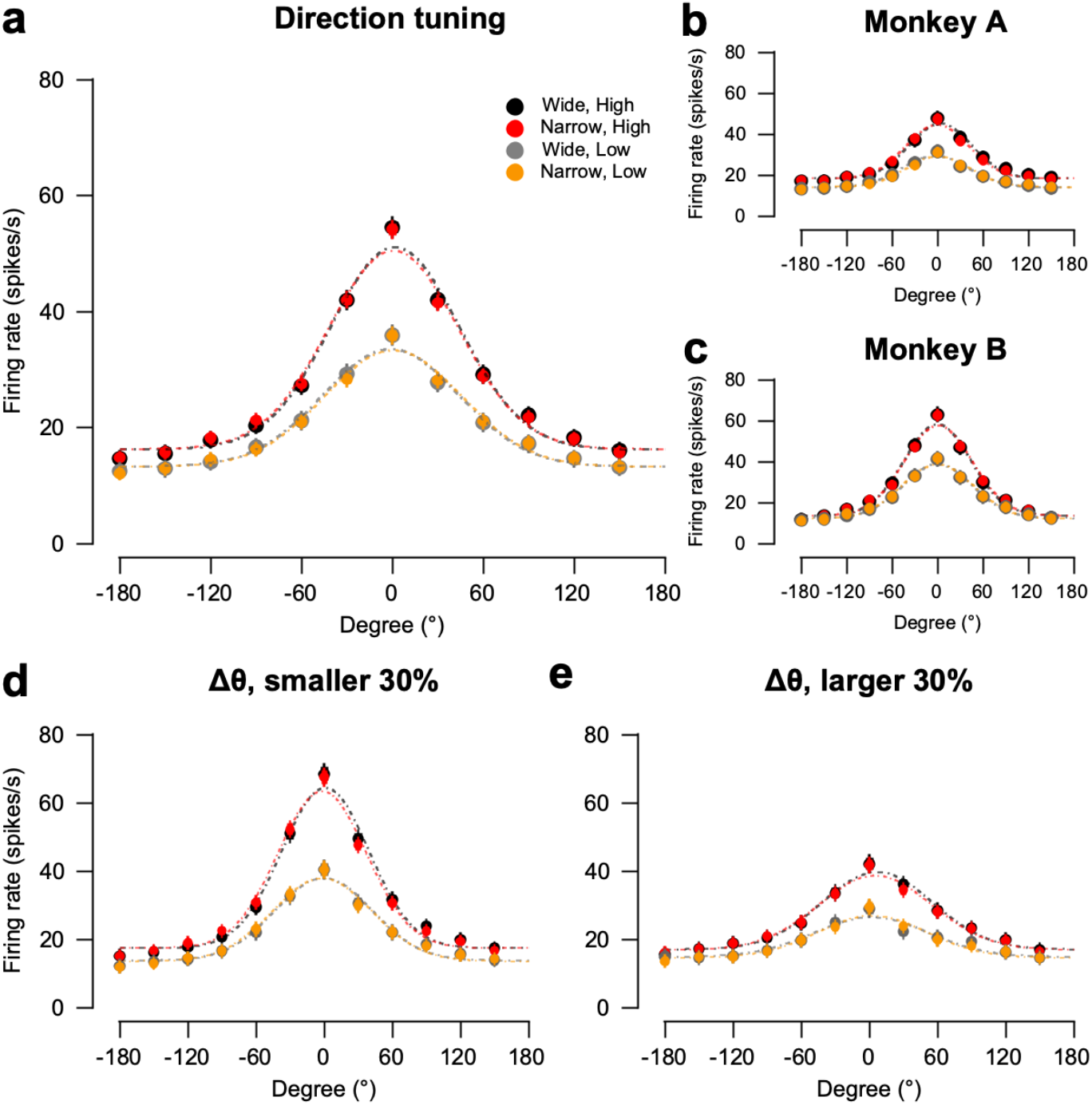
Direction tuning responses of MT neurons in a passive fixation task. **a** Mean direction tuning across all MT neurons in the combined data from the two monkeys. The direction tuning responses of individual neurons are realigned relative to each neuron’s preferred directions (filled circles) and fitted to a Gaussian function (dash-dotted lines). **b, c** Mean direction tuning of MT neurons in each monkey. **d, e** Mean direction tuning of top and bottom 30 % neurons based on the size of the difference between each neuron’s preferred direction and prior direction in the combined dataset.

To further test whether the change in the direction tuning function of MT neurons occurred only when the preferred direction of each neuron was far from the prior direction, we divided the neurons into two groups; one with larger direction discrepancies between each neuron’s preferred direction and the prior direction, and the other with smaller discrepancies (i.e., top and bottom 30 % of neurons based on the direction discrepancies). We then averaged the direction tunings of the neurons in each group. As shown in the results from the entire population, the direction tunings of the neurons in both groups were not different between the wide and narrow prior blocks (Figs. 4d, e, the height of Gaussian function in the wide and narrow prior blocks; bottom 30 % group of 54 for high contrast: 64.33 and 63.36, bottom 30 % group of 53 for low contrast: 37.89 and 37.86, top 30 % group of 54 for high contrast: 39.68 and 38.68, top 30 % group of 53 for low contrast: 26.44 and 26.86). This suggests that the systematic response reduction did not result from the passive neural response changes induced by the repetitive presentation of the motion stimulus. These results exclude adaptation accounts and demonstrate that the systematic reduction in neuronal response only appears when prior knowledge of motion direction is used.

### Prior expectation is represented in the neural subspaces

The monkeys would switch the usage of prior expectation across blocks of trials; therefore, in the narrow prior block, the expectation-related signal could modulate the activity of area MT neurons even before the pursuit target appearance. When this possibility was tested in the average PSTHs during the pursuit task (and fixation task), prior expectations did not modulate the preparatory activities (Figs. 2a, b for both monkeys, Figs. 5a, d for Monkey A and Supplementary Figs. 3a, d for Monkey B). The mean firing rate of the MT neurons differed only between the wide and narrow prior blocks after pursuit target onset (Figs. 2a, b). Although there was no effect of prior expectations on average preparatory neural activity, it may be present in the activity pattern of the neural population. Recent work has shown that context, sensory inputs, and perceptual choices can be represented in task-relevant dimensions of population responses as well as in single-unit responses in the prefrontal cortex ^42,43^. To identify whether the MT population encodes the task variables, including prior expectation and sensory information, we used the targeted dimensionality reduction (TDR) method ^42^. As only the pursuit targets of the prior directions were presented in both the wide and narrow prior blocks, we restricted subsequent analyses to the neural responses to the prior directions (see Supplementary Fig. 4 for the analysis that included responses to all stimulus directions). For the population analysis, we constructed population responses by pooling all the units that were recorded primarily in separate sessions. First, we estimated linear regression coefficients for the prior expectation and stimulus contrast, respectively. To de-noise the regression coefficients, we projected them into the subspace spanned by dominant 12 principal components of population activity and then reprojected the denoised coefficients in the orginal basis. The regression coefficients for the prior, showing dependencies of the MT neural responses on the prior block, were temporally correlated during the preparatory period (Fig. 5b for Monkey A and Supplementary Fig. 3b for Monkey B, see Supplementary Figs. 4a, b for the regressions coefficients for the contrast). This temporal correlation suggests that the MT population activity retains prior information, even before the appearance of the pursuit targets. Next, we defined task-related axes by time averaging the regression coefficients for prior and contrast and then orthogonalizing them. The averaged population responses across trials were projected onto these axes, generating neural trajectories in the task-related subspace (see Methods for details). Neural trajectories show the temporal evolution of the average population response during the pursuit task trial. The projected population response moved in the direction of high contrast (upward) and wide prior (rightward) after target onset, driven by sensory information from the stimulus (Figs. 5c, f and Supplementary Figs. 3c, f). The general direction of these trajectory changes is due to the sensory inputs: sensory stimuli have luminance contrast information, which has a range of 0-100 % (upward changes), and the appearance of the sensory stimulus will reduce the need for the specific expectations about the stimulus (rightward changes). Notably, the population responses in the wide and narrow prior blocks were separable in the prior-axis (Fig. 5c and Supplementary Fig. 3c for Monkey A and B, respectively), where clear separation was already observed at the preparatory activity (Insets in Fig. 5c and Supplementary Fig. 3c). We quantified this separation by estimating the distance of neural trajectories on the prior-axis between the two prior blocks. The subspace distance was roughly maintained throughout the trial including the preparatory period in the pursuit task (Fig. 5h for Monkey A and Supplementary Fig. 3h for Monkey B, the mean difference of prior-axis projected population responses between the wide and narrow prior blocks during the pursuit task was 1.04 and 1.18 in Monkey A; 0.60 and 0.72 in Monkey B for high and low contrast cases, and that during the preparatory period was 0.42 and 0.60 in Monkey A; 0.48 and 0.35 in Monkey B for high and low contrast cases, two-sided permutation test, p < 0.05, see Methods for details).

**Fig. 5:**
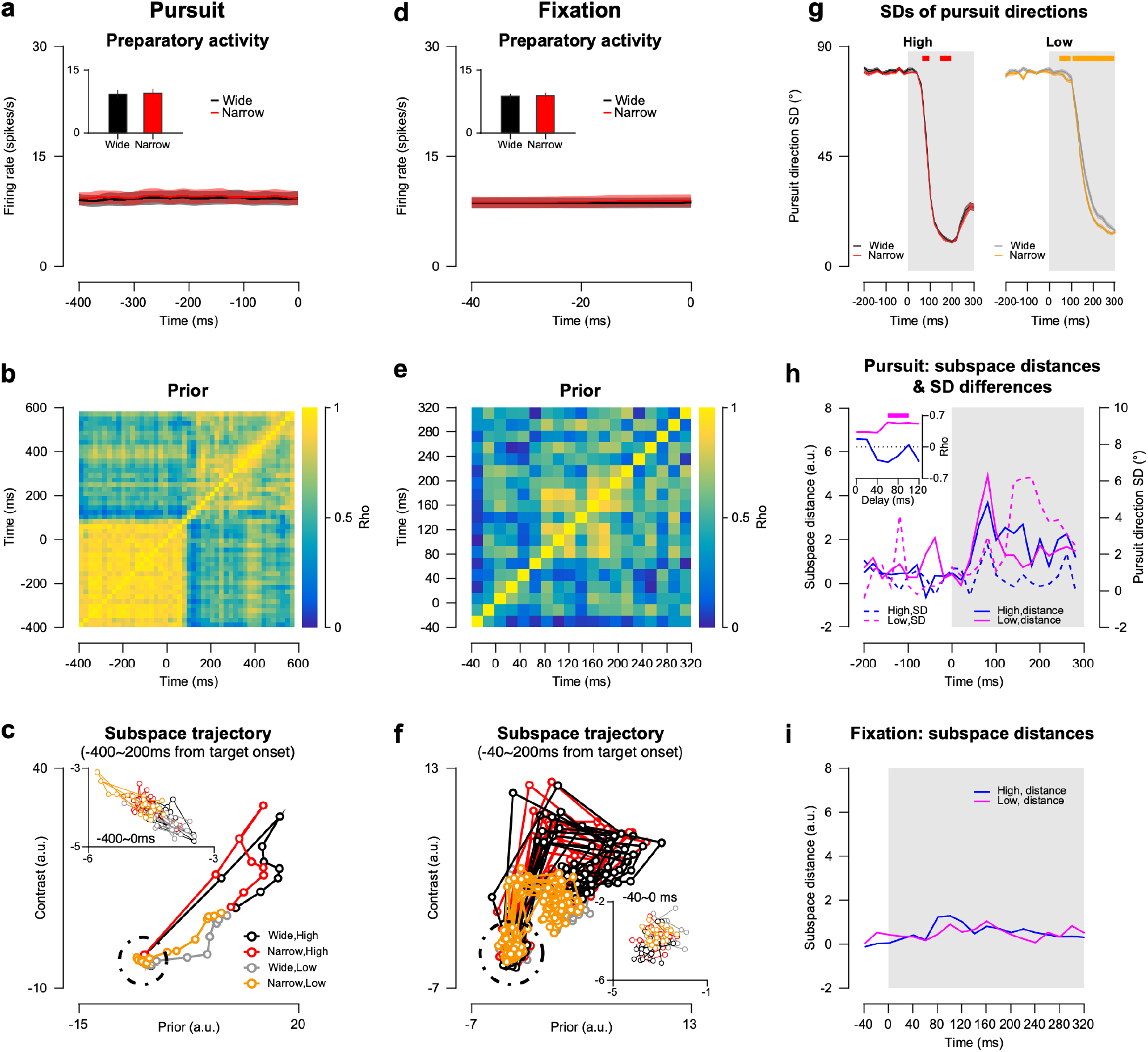
Latent dynamics of MT population responses during pursuit and fixation task. **a**,**d** PSTHs relative to the target onset and mean preparatory activities of the recorded MT neurons (inset), in each prior block during the pursuit task (**a**) and fixation task (**d**). **b**,**e** Autocorrelation matrix of regression coefficients for prior during the pursuit task (**b**) and fixation task (**e**). **c, f** Temporal trajectories of average population responses in the task-related subspace consisting of prior and contrast axes during the pursuit task (**c**) and fixation task (**f**). Trajectories of preparatory activity(–400ms-0ms in the pursuit task and –40ms-0ms in the fixation task) are shown in the inset plots. **g** SDs of pursuit directions with high (left) and low (right) contrast stimuli as a function of time relative to the target onset. The red and yellow thick lines indicate that the SD differences between the two blocks are significant (cluster-based permutation test, p <0.05). **h** Distances in population responses projected on the prior-axis of the subspace between the wide and narrow prior blocks as a function of time (solid lines), and the differences of pursuit direction SDs between the two priors as a function of time (dotted lines). The blue and magenta lines indicate the high contrast and low contrast conditions, respectively. Temporal coherences between subspace distances and SD differences with different latencies of SD differences from subspace distances are shown in the inset plots. The thick top line in the inset indicates that the FDR-corrected p-values are less than 0.05. **i** Distances in the prior-axis projected population responses between the two prior blocks during the fixation task.

Latent dynamics along the prior-axis had functional relevance to pursuit behaviors, consistent with the simulation results. We calculated the SDs of the pursuit directions as a function of time to obtain the dynamics of behavioral variability during the pursuit trial (Fig. 5g for Monkey A and Supplementary Fig. 3g for Monkey B). The SDs rapidly decreased during the open-loop period (approximately 100-200ms from the stimulus onset). The reduction in SD in the narrow prior block was stronger than that in the wide prior block, particularly when the contrast of the sensory stimulus was low. To determine whether the subspace distance is related to behavioral variance, we compared the temporal change in neural distances with that in pursuit SD differences. Both the subspace distances and SD differences were the highest shortly after the pursuit target onset (Fig. 5h for Monkey A and Supplementary Fig. 3h for Monkey B). Only in the low contrast condition, the subspace distance showed a high temporal correlation with the SD differences, and the subspace distances always lead the SD differences (significant correlation of 0.5 between the two indicates that SD differences appeared 60-120ms later than subspace distances in Monkey A; 40-80ms later in Monkey B, Insets in Fig. 5h and Supplementary Fig. 3h, fDR corrected, alpha = 0.05). This result additionally suggests that the neural modulation of MT population responses by prior expectations may contribute to reducing the variability in oculomotor behavior.

Neural subspace modulation was specific to the pursuit task. When the same population analysis was applied to the neural population data from the fixation task, the modulation of the population neural spontaneous activity along the prior-axis of the subspace disappeared. Regression coefficients for the prior were not correlated across time for almost the entire duration (Fig. 5e for Monkey A and Supplementary Fig. 3e for Monkey B). Furthermore, the trajectories within the neural subspace were not separable between the wide and narrow prior blocks before visual stimulus appeared, thus demonstrating the absence of prior expectation-related input to the area MT (Inset in Fig. 5f for Monkey A and Inset in Supplementary Fig. 3f for Monkey B). There was no difference in neural trajectories projected to the prior-axis between the wide and narrow prior blocks during the fixation task (two-sided permutation test, p > 0.5). These results suggest that the expectation signal was present in the neural subspace of MT activity only when the monkeys were engaged in using prior knowledge in the task.

## Discussion

In this study, prior expectations reduced the variation in pursuit directions and discriminately reduced the responses of MT neurons, only when the sensory evidence was weak, depending on their preferred directions. Using data-based simulations and multiple decoders (SVM, PVD, and MLE), we demonstrated that the systematic response reduction improves the neural representation of motion direction by sharpening population tuning in area MT, which accounts for the behavioral improvement. Additionally, TDR analysis of the population neural responses demonstrated the existence of the prior expectation signal in the neural subspace of spontaneous activity and the functional relevance of the subspace dynamics to the behavioral variability reduction observed in pursuit initiation.

### Neural origins of prior expectation

The Bayesian observer model has successfully revealed the integration of prior expectation and sensory information in smooth pursuit eye movements ^35^; however, the corresponding neural areas and mechanisms for this inference are largely unknown. Notably, whether the sensory area only represents the likelihood function of visual input or also represents the prior information and the posterior probability distribution is unknown. The hierarchical Bayesian inference framework for visual processing provides a theoretical ground for understanding the neural mechanisms underlying the interplay between sensory and top-down signals ^44–47^. Within this framework, higher-order brain regions may suppress lower-order neural responses, particularly when they are “incongruent” with prior expectations ^46^. Thus, bottom-up sensory signals are reduced in such a manner as to increase the signal-to-noise ratio. Recent human neuroimaging studies have provided empirical support for this hypothesis by showing expectation-driven modulation of neural activity and representation in the visual cortex ^29,31,32^. However, these neuroimaging data cannot explain how the visual system improves the sensory representation by silencing the incongruent responses; additionally, the neural responses that are incongruent with the expectations are ambiguous. Here, we propose a neural substrate of prior expectation in the visual cortex that the neural responses reduce to sharpen population direction tuning. The simulation revealed that prior expectations can change pursuit direction bias and variability via the systematic reduction in MT neural responses. These results underpin the Bayesian brain hypothesis by showing that the sensory area may represent not only the likelihood function but also the posterior distribution to which prior information is applied.

A possible alternative framework for sensory processing is the traditional feedforward model. Under this framework, the integration of sensory evidence and prior information occur in the parietal and frontal areas but not in the sensory areas ^48,49^. A neurophysiological study by Rao et al. ^34^ supported this perspective; the responses of neurons in the macaque lateral intraparietal area were affected by prior expectations of stimulus motion, whereas almost no change was observed in MT neuronal responses during a direction discrimination task using saccadic eye movements. However, our results of the reduced MT responses by prior expectation contrast these findings. At least two possible factors could attribute to the different effects of prior expectations on area MT neurons. First, the previous research and this study used different eye movement systems (saccades and pursuits) for behavioral tasks. Although saccadic and pursuit eye movements share some common parts of the oculomotor network, they have largely disparate phylogenies and functions ^50^. Therefore, although area MT is involved in both saccades and pursuits, the influence of cognitive modulation on MT neural activity may differ depending on the eye movement system. Second, the two studies used different behavioral paradigms to control the monkeys’ prior expectations of the motion direction of the stimuli. Rao et al. used an arrow cue for “transient” top-down modulation, whereas we used “cumulative” learning based on experience-dependent prior knowledge from previous trials. These different strategies for developing an expectation of motion direction may have resulted in different neural mechanisms.

In our results, the prior expectation signal was present in single neuronal responses in area MT only after the pursuit target appeared but was also represented as the pattern of population responses even before the pursuit target onset. TDR analysis ^42,51^ clearly showed the effect of prior expectation on spontaneous population activity (Figs. 5b, c for Monkey A, Supplementary Figs. 3b, c for Monkey B), and this effect was task-specific (absence of the effect in the fixation task, Figs. 5e, f for monkey A, Supplementary Figs. 3e, f for monkey B). These results strongly suggest the existence of the ‘prior’ inputs in the activity of MT neurons. Notably, the prior expectation signal that appeared in the spontaneous activity was maintained during smooth pursuit initiation, regardless of the strength of sensory inputs (neural subspace distance between the wide and narrow prior blocks: Figs. 5c, h for Monkey A, Supplementary Figs. 3c, h for Monkey B). Neurons in area MT may receive top-down expectation-related inputs from other brain regions, and these signals may be dynamically integrated with feedforward sensory inputs evoked by the pursuit target. Therefore, the neural activity in area MT may represent the posterior probability distribution by integrating the prior inputs and sensory inputs during the presentation of the pursuit target. Both univariate (unit) and multivariate (population) analyses suggest the reliability-weighted integration of these two inputs by showing that the expectation-evoked neural changes accounted for the behavioral modulation in pursuit initiation only when the sensory evidence is weak (Figs. 3 d-f and Fig. 5 h). However, because most neurons were not simultaneously recorded, our analyses could not directly show the modulation of both the population tuning and population response pattern by prior expectations. Future work should aim to identify the contribution of population-level representations of sensory and cognitive information to behavior through direct analysis of the trial-by-trial relationship between population activity and behavior.

Which brain regions provide this task-specific expectation signal to area MT? A recent study ^14^ demonstrated that FEF_SEM_ represents the prior expectation of speed through preparatory activity; prior expectation of faster speed results in higher spontaneous activity during fixation before visual motion onset. If the preparatory activity in FEF_SEM_ also represents the prior expectation of motion direction, the source of prior information could be FEF_SEM_, although the manner of representing the directional expectation might be different from simple gain modulation. In addition, the precision-weighted integration between the prior and likelihood functions of motion speed was represented in the evoked responses of FEF_SEM_. Because both area MT and FEF_SEM_ represented prior-likelihood integration, the two sites appear to have an essential role in the effect of prior expectation on smooth pursuit eye movements while interacting with each other. A previous study reported that prior expectations of the direction of a visual stimulus increase the functional connectivity between area MT and prefrontal cortex ^52^. Investigating the role of each region and the interaction between the two in implementing Bayesian inference during perceptual decision-making and sensorimotor behavior is an important direction for future research.

### Expectation and Attention

Expectation and attention have a similar facilitatory effect on visual perception: expected and attended stimuli are more easily detected and recognized than unexpected or unattended stimuli. Therefore, the effects of expectation on perceptual decisions and sensorimotor behavior have often coalesced with those of attention in the empirical literature. In numerous cases, changes in perception and behavior due to expectation can also be explained by attention, and the distinction between these effects is unclear. For example, the behavioral and neural effects of expected stimuli might be caused by decreased attention and vice versa. Although the mechanisms overlap and interact, neither expectation nor attention is solely responsible for the influences of the two on visual perception. For fully understanding expectation and attention, they must be conceptually dissociated ^53–56^. Although expectation and attention were not strictly separated in our task design, the behavioral results were not due to spatial attention as the pursuit targets were always presented at the same position (fovea) between the two prior blocks. Unlike spatial attention, feature-based attention (i.e., attending to the direction of motion) affecting the perceptual behavior during our task might be concerning. Determining whether the results were from prior expectation or feature-based attention from behavioral data alone is challenging; however, the neural results may provide insight into this.

In terms of neural modulation, attention increases the responsivity of sensory neurons ^57^, whereas expectations decrease the neural activity in sensory areas ^29,31,58^. In area MT, spatial ^59,60^ and feature-based attention ^61,62^ increase the gain of the direction tuning curves of MT neurons without changing the tuning bandwidths. However, the effects of expectation on these neurons remain largely unknown. In this study, prior expectation reduced the visual responses of MT neurons, which is consistent with the results of previous neuroimaging studies, and importantly, the reduction in neural activity might contribute to improving behavioral reliability. This supports the observation that the reduced responses were not due to lower attention but a higher expectation because less attention to stimuli cannot enhance behavioral performance.

Furthermore, previous attention studies reported that behavioral performance can be improved by reducing the trial-by-trial variability of single neuronal responses in the visual cortex ^63–65^. Thus, if prior expectation and attention operate on a shared mechanism, prior expectations may reduce the variation in eye movements by modulating spike count variability of MT neuronal responses. We measured the mean-normalized variance (i.e. Fano factor) of spike counts and compared it across different conditions. A transient decrease in the Fano factor occurred after stimulus onset in all conditions (wide/narrow prior × high/low contrast) and the decrease was larger in the high contrast cases than in the low contrast cases (Supplementary Figs. 6a, b), which was consistent with previous results ^66^. However, the Fano factor did not differ between prior conditions. In Monkey A, when the stimulus contrast was high, there was a period in which the Fano factor in the narrow prior block was lower than that in the wide prior block (cluster-based permutation test, *p* < 0.05). However, this effect was observed after the eye began to move; therefore, it might be caused by the difference in eye movements. Additionally, this effect was negligible in Monkey B. These results suggest that the prior expectation of motion direction does not significantly change the variability of single-neuronal spiking.

In addition to single neuronal variability, the modulation of the trial-by-trial spike count correlation between neurons can affect perceptual performance. Previous studies demonstrated that the attention-induced reduction in the inter-neuronal correlation has a tight relationship with enhancing behavioral performance ^63,65,67,68^. To test if prior expectations modulate the inter-neuronal correlation, we measured and compared the trial-by-trial correlations between pairs of neurons across the wide and narrow prior blocks. However, the pairwise spike count correlations between neurons did not differ between prior conditions (Supplementary Figs. 6c–f). These results suggest that, unlike the attentional modulation in the neural variability, neither the trial-by-trial spike count variability of single neurons nor trial-by-trial correlation between pairs of neurons can explain the reduction in the behavioral variability by prior expectations. Therefore, the neural mechanisms of expectation and attention may differ during visual perception and oculomotor responses. A well-designed experimental paradigm would be necessary to properly dissociate expectation effects from attention effects and systemically explore the similarities and differences between the two mechanisms.

### Expectation and Neural adaptation

Previous studies have reported the adaptation of MT neurons to the repeated motion direction of visual stimuli ^41,69–71^; neural adaptation reduced the amplitude and width of the direction tuning curves of MT neurons, and the effects were the largest when the preferred direction of the MT neuron and the direction of the adapting stimulus matched. In this study, pursuit targets were presented for the prior direction twice as often as for the other directions in the narrow prior block. However, although the neuronal responses to the prior direction were reduced, this reduction differed from the passive response modulation of neural adaptation for the following reasons. First, the direction tuning curves of MT neurons in the narrow prior block were similar to those in the wide prior block when the animals passively fixed their eye gazes at the central fixation point (Fig. 4). Second, contrary to what was expected from neural adaptation, the effect was stronger when the prior direction differed from the preferred direction. In addition, the amount of response reduction by prior expectation did not depend on the extent to which the receptive field and pursuit target overlapped (Supplementary Fig. 5). These differences imply that the reduced responses of MT neurons in this study may not have been due to prolonged exposure to repeated visual motion but rather because of cognitive feedback signaling, which is different from the simple mechanistic reaction of the neurons to adjust constant sensory input.

## Methods

Two adult male rhesus monkeys (Macaca mulatta) weighing 9–11 kg were used for the neurophysiological experiments. All research protocols were approved by the Sungkyunkwan University Institutional Animal Care and Use Committee. Before the experiments, we performed two separate surgeries. A head holder was implanted on the skull for head restraint, and after which a cylindrical chamber fabricated from PEEK was implanted on the skull close to the lunate sulcus for an angled approach to area MT. During the surgeries, each monkey was under isoflurane anesthesia, while antibiotics and analgesics were administered postoperatively to minimize infection and pain.

### Visual stimuli and behavioral paradigm

Visual stimuli were presented on a gamma-corrected 24″ CRT monitor (HP1230, 1600 × 1200 pixels, 85 Hz vertical refresh rate). The monitor was placed 570 mm from the animal, and it covered 38.67° by 29.49° of the horizontal and vertical visual field, respectively. The background on which visual stimuli were presented was gray with a luminance level of 36.8 cd/m^2^ (luminance range was 0-73.68 cd/m^2^). The presentation of visual stimuli and recording of eye movement data were controlled using a real-time data acquisition system (Maestro version 3.3.11). A custom-built photodiode system was used to ensure the accurate timing of the visual stimuli.

The two monkeys were trained in a smooth pursuit eye movement task (Fig. 1a). The pursuit target was a random dot kinematogram, and its size in each day’s experiment was determined as one of 4° × 4°, 8° × 8°, or 12° × 12° depending on the location and size of the receptive fields of the recorded MT neurons. Each trial began when the animals fixated their eyes on a small dot (fixation spot) at the center of the monitor screen. After a randomized fixation duration (800, 1300, or 1800 ms), the dot patch appeared at the center of the screen or 1°–2° displaced from the center to the opposite direction of the target direction. Subsequently, a local motion, whereby all the dots inside the invisible circular window moved in the target direction at a given speed but the invisible window did not move, was implemented. Following the local motion, both the dots and windows moved together at the same speed and in the same direction as the local motion for 500–700 ms. If the animals maintained their eyes at the center of the pursuit target within a 4° window during the target movements, they were rewarded with a drop of water or juice.

To control the animals’ prior expectation of the motion direction of the pursuit target, we used two types of blocks: wide and narrow prior blocks (Fig. 1b). In the wide prior block, the fixation spot was red and the pursuit target direction was randomly and evenly selected from three directions, which were 120° apart from each other. Therefore, the animals’ expectations for the incoming motion direction would be widely distributed across the directions. Conversely, in the narrow prior block, the fixation spot was green and the direction of the pursuit target was one of the three narrowly distributed directions (a central direction and ± 15° of the direction). The central direction was presented twice as often as the other two directions. Therefore, the animals tended to develop a prior expectation for this central motion direction or the expectation would be narrowly distributed around the central direction. Therefore, we termed this direction the “prior direction.” The prior direction was set by considering the preferred directions of the recorded neurons. Notably, the identical prior directions were included in both the wide and narrow prior blocks to compare the effect of prior expectations on behavioral and neural responses under the identical sensory stimuli. The narrow and wide prior blocks consisted of 252 and 378 trials, respectively. The first block in each day’s experiment was randomly selected and alternated between the two prior blocks. We typically collected four blocks for each category, resulting in 2520 trials in total.

Further, the precision of the sensory evidence (sensory stimulus) was controlled by randomly selecting a pursuit target stimulus from the two stimulus types in each trial: one was a random-dot kinematogram with 100 % luminance contrast (high contrast) and the other was a random-dot kinematogram with 12 % or 8 % (for Monkey A and B, respectively) luminance contrast (low contrast). To reduce the precision of motion direction further in the low contrast case, a random-walk direction noise ^72^ was included in the stimulus; the directions of all dots were randomly selected from uniform distribution bounded by ±60° of the target direction (Fig. 1c).

### Data collection and analysis

Horizontal and vertical eye positions were separately recorded at a sampling rate of 1 kHz using an infrared video tracking system (EyeLink 1000 Plus, SR Research Ltd). A low-pass Butterworth filter with an order of two and cutoff frequency of 20 Hz was applied on the horizontal and vertical components of the eye position. Subsequently, the filtered eye positions were differentiated to obtain the horizontal and vertical eye velocities. To focus on the initiation of the smooth pursuit eye movements and remove the influence of saccadic eye movements (saccades) on the behavioral and neural responses, trials with saccades that occured in a time window between –100 and 250 ms were removed from the onset of the pursuit target. Next, the eye velocity traces were rotated such that their average direction was 45°. The rotated eye velocity during the open-loop period (100 ms from pursuit onset ^36^) was decomposed by using horizontal and vertical templates ^73,74^, which were the average of each rotated eye velocity component between –20 and 100 ms from the average pursuit latency (mean pursuit onset for high and low contrast stimuli: 76 ms and 123 ms in Monkey A; 57 ms and 105 ms in Monkey B). The optimal pursuit latency and scaling factors were estimated through the sliding and scaling of the two templates with the least-squares method and NOMAD algorithm ^75^.

A single electrode or an 8-channel laminar probe (Multitrode Type I, Thomas RECORDING) with impedances from 0.5 to 1 MΩ (at 1 kHz) was used to record spikes and local field potentials (LFPs) in the MT using the Motorized Electrode Manipulator (MEM) system (Thomas RECORDING). Extracellular electrical signals from the electrode were high-pass filtered at a cutoff frequency of 150 Hz, digitized at a sampling rate of 40 kHz for action potentials, and low-pass filtered at a cutoff frequency of 170 Hz, and digitized at a sampling rate of 1 kHz for LFPs (OmniPlex). The approximate receptive field location and size of a well-isolated MT neuron were measured using a hand-controlled visual stimulus while the monkeys fixated their eye gaze on a stationary spot at the center of the monitor. The MT neurons whose receptive fields were near the fovea were analyzed as the pursuit targets were presented at or near the fovea.

To isolate single-unit responses, we sorted the spikes offline (Offline Sorter, Plexon) using principal component analysis and the waveforms of recorded neural activities. For further analysis, we included only the neurons whose spikes formed distinct clusters in the principal component space. Additionally, we checked and removed sorting errors by inspecting time stamps of the sorted spikes with a time resolution of 1 ms. Thus, we obtained 137 neurons from 59 days of recordings of Monkey A and 120 neurons from 67 days of recordings of Monkey B. For behavioral analysis, only days with more than 50 trials for each condition (four conditions: wide/narrow prior block and high/low contrast stimuli) were used (59 days for Monkey A and 65 days for Monkey B).

### Spike count analysis

The responses of MT neurons to the prior direction stimulus between the wide and narrow prior blocks were compared to probe the effect of prior expectations on the neural responses. This was conducted by constructing a PSTH for each prior condition and smoothing it using a 10 ms SD Gaussian filter (Figs. 2a, b). Additionally, the spikes in the 100 ms duration time window from the spike latency of each neuron were counted and compared between the prior blocks (Insets in Figs. 2a, b, and Fig. 2c). To measure each neuron’s spike latency, the spiking responses across all repetitions of the given stimulus contrast (high or low) were averaged over time, and the latency was estimated by visual inspections of the mean PSTH. If the neuronal latency could not be visually determined because of the lack of distinct stimulus-related responses, the neuron was excluded from the analysis. Thus, in monkey A, 136 and 111 neurons were used for the high and low contrast cases, respectively, and in monkey B, 117 and 110 neurons were used for each contrast case. To calculate the mean-normalized variance (variance/mean; Fano factor) of spike counts as a function of time, a 100 ms (±50 ms) sliding time window with 20ms steps was used (Supplementary Figs. 6a, b). For the in silico simulations, the Fano factor estimated from the time period between 0 and 200 ms after the pursuit target onset was imposed (Fig. 3 and Supplementary Fig. 2). To estimate the correlation between the difference in the prior and preferred directions and the firing rate ratio of the narrow prior block to the wide prior block over time, the Spearman’s correlation coefficient was computed for each 60 ms (±30 ms) sliding time window from pursuit target onset to 180 ms with 5 ms time intervals (Figs. 2d–i). If the neuronal latency or preferred direction was unclear, the neuron was excluded from this analysis. Thus, 95 and 75 neurons from Monkey A were used for the high and low contrast cases, respectively, and 86 and 69 neurons from Monkey B were used for each contrast cases.

### Direction tuning measurement and estimation

Before the smooth pursuit eye movement task, the direction and speed tuning of the isolated MT neurons were estimated in a separate direction tuning task. To assess the direction tuning of the neurons, stimulus motion was presented in 12 directions (0°–330° in 30° increments) at the preferred speed of the MT neurons while animals fixated their eye gaze on a central stationary spot. A circular patch of random dots with 100 % coherence served as the visual stimulus, and the location and size of the visual stimulus were set according to the cells’ receptive field properties. Each trial began with a fixation duration between 600 and 1200 ms, and then six different motion stimuli were presented for 256 ms with 300 ms of interposed stationary epochs. For the direction tuning curve, the spike counts for 256 ms, after 50 ms from the stimulus onsets, were fitted to a Gaussian or circular Gaussian function:

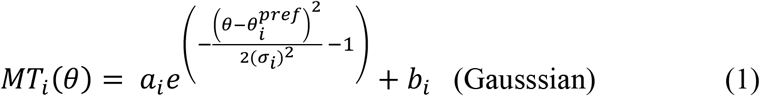

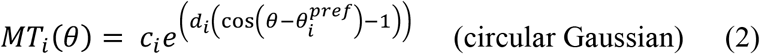

where *θ* is the 12 motion directions of the visual stimulus; *a*_*i*_, *b*_*i*_, 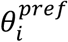, and *σ*_*i*_ are the parameters of the Gaussian function of *i*th MT neuron, where 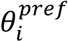 and *σ*_*i*_ denote the mean and SD of the Gaussian function, respectively; *c*_*i*_ and *d*_*i*_ are the parameters of the circular Gaussian function of *i*th MT neuron, and 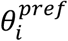 indicates the preferred direction of the MT neuron. The direction tuning responses for the different luminance contrasts were obtained by randomly interleaving the high contrast (100 % luminance contrast for both monkeys) and low contrast (luminance contrast of 12 % for Monkey A and 8 % for Monkey B) patches as the visual stimuli. For speed tuning of the neurons, the motions at nine speeds (0, 1, 2, 4, 8, 16, 32, 64, and 128 °/s) with the preferred direction of the neurons were presented. The speed tuning responses were fitted to a log Gaussian function ^76^. If the direction tuning exhibited a bimodal shape or the estimated preferred speed was extremely high or low, the orientation tuning was also quantified to identify whether the neuron had an orientation selectivity rather than direction selectivity. Six orientations (0°, 30°, 60°, 90°, 120°, 150°) were used for the orientation tuning task and excluded the neurons when its orientation tuning responses were more robust than the direction tuning responses.

Additionally, the direction tuning responses of MT neurons in the middle of the smooth pursuit task were intermittently measured by randomly inserting trials of the direction tuning task. The visual stimuli for the direction tuning measurement were identical to the pursuit targets, but they were placed and remained in the center of the receptive fields to evoke strong MT neuronal responses. To characterize the tuning properties, the directional responses of each neuron measured during the pursuit task was fitted to a Gaussian or circular Gaussian function based on their explained variance, except that only a circular Gaussian function was used to estimate the direction tuning functions of the MT neurons for the simulation. We excluded neurons from the further analysis if the fitted tuning curves explained less than 50 % of the response variance or if the tuning measurement in either prior block was missing. Accordingly, 82 and 86 neurons from Monkey A were used for the high and low contrast cases, respectively, and 83 and 80 neurons from Monkey B were used for each contrast case to quantify the direction tuning measured during the smooth pursuit task. To obtain the average tuning responses of the neurons, we realigned the direction tuning responses of individual neurons relative to each neuron’s preferred direction such that all preferred directions were 0° (Fig. 4). The preferred direction of each neuron was determined based on the tuning function for the high contrast stimulus to obtain more stable tuning parameters. The width of the tuning function was estimated by calculating its full width at half maximum of the tuning function.

### Inter-neuronal spike count correlation

Trial-by-trial spike count correlations between pairs of MT neurons were calculated using Spearman’s correlation coefficient (Supplementary Figs. 6c–f). We used 100 ms (±50 ms) sliding time windows from –200 to 400 ms after the pursuit target onset with a step size of 20 ms. Pairs of neurons were included in the sample only if the number of trials for the prior direction was more than 50 in both the wide and narrow prior blocks. This criterion was applied separately for each stimulus contrast condition. This resulted in the analysis of 165 (high contrast) and 99 (low contrast) pairs from Monkey A and 123 (high contrast) and 117 (low contrast) pairs from Monkey B.

### Simulating and decoding of MT population responses

For the in silico simulations, the firing rates of MT neurons in response to stimulus motion were modeled as a circular Gaussian function (Eq. 2) based on our experimental dataset ^69^. Each circular Gaussian function parameter for the responses of *i*th model MT neuron (*c*′_*i*_ and *d*′_*i*_) was randomly selected from a gamma distribution fitted to the estimated parameters of the direction tuning functions of the recorded MT neurons using the *gamfit* function in MATLAB:

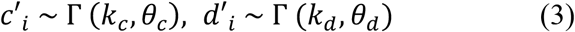

where *k*_*c*_ is the shape parameter; *θ*_*c*_ is the scale parameter of the gamma distribution of *c*; *k*_*d*_ and *θ*_(_ are the shape and scale parameters of the gamma distribution of *d*, respectively. MT neuronal responses to the high and low contrast stimuli were simulated (*k*_*c*_= 1.74 and 1.28; *θ*_*c*_ = 36.83 and 31.16; *k*_(_ = 1.30 and 1.18; *θ*_(_ = 1.19 and 0.73 for the high and low contrast stimuli, respectively). Preferred directions 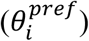 ranged from –179° to 180° in 1° increment, and each preferred direction was assigned to 10 simulated neurons. Thus, the model totally included 3600 (360 × 10) neurons for every simulation. Trial-by-trial variation in individual neuronal responses and trial-by-trial correlation between neurons were simulated using the mean Fano factor and mean inter-neuronal correlation over 200 ms from the pursuit target onset estimated from the MT recordings. Because we could not find any evidence of significant changes in Fano factors and inter-neuronal correlations across prior conditions, we averaged Fano factors and inter-neuronal correlation coefficients from the wide and narrow prior blocks in each contrast case. (Fano factor for high contrast: 1.11, low contrast: 1.37; inter-neuronal correlation for high contrast: 0.02, low contrast: 0.06). The correlation between neuronal responses was simulated using the Cholesky decomposition method with a constant inter-neuronal correlation coefficient for each contrast condition ^77^. To model the reduction effect of prior expectation on neuronal responses, we multiplied the simulated MT responses by the exponential of the regression coefficient between Δθ (preferred direction–prior direction) and the log firing rate ratio (narrow/wide prior) computed from the experimental data (regression slope at 85 ± 30 ms from the pursuit target onset for high contrast, –0.0003, and for low contrast, –0.0025, see Figs. 2g, h).

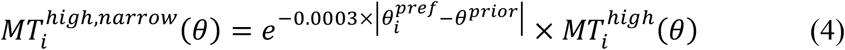

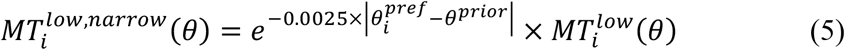

where *θ*is the direction of target motion; 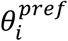 is the preferred direction of the i_th_ neuron; *θ*^*prior*^ is the expected direction (the prior direction); and 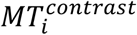 is the average firing rate of the i_th_ neuron to the target direction in each contrast, from Eq. 1.

### Support Vector Machine

The SVM classifiers were trained with 80 % of the model MT population to discriminate two motion directions using the *fitclinear* function in MATLAB. With the remaining 20 % of the population neural activity, the trained classifier predicted one of two directions, which were 0° and a direction between 0° and 10° (step sizes of 0.5°). We simulated 200 trials of neuronal population responses to each motion direction and repeated each pair of simulations 100 times to validate the predictive performance of the SVM classifier.

### Population Vector Decoder

Population vector averaging was used to estimate the direction information represented in the neuronal population activity. Using the simulated MT neuronal responses, the PVD predicted the direction of the stimulus motion as follows:

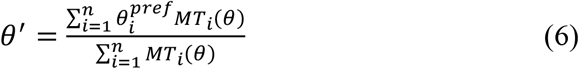

where *θ* is the direction of motion of the visual stimulus; 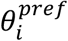 is the preferred direction of *i*th model MT neuron; MT_i_(*θ*) is the response of the *i*th neuron obtained from Eq. 1. The target direction *θ* was set to 0° for the prior direction (Fig. 3e) or ±15° for the outer directions (Supplementary Fig. 2c, d). To compensate for the estimation error resulting from asymmetries in contributing neural population in PVD, the preferred directions of model MT neurons ranged from –194° to 165° or from –164° to 195° (instead of the ranging from –179° to 180°) when the target direction was –15° or 15°, respectively. We simulated 200 trials and computed the SD of the predicted directions *(θ*^*′*^) in each simulation. This procedure was repeated 100 times.

### Maximum Likelihood Estimation

The simulated MT responses were also decoded using maximum likelihood estimation ^38–40^. The log likelihood of a motion direction *θx* can be expressed as:

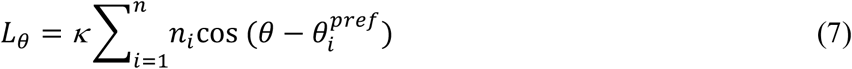

where *K* is the concentration parameter of the circular Gaussian function, which is the tuning function of the neurons; *n*_*i*_ is the spike counts of *i*th MT neuron in response to the direction *θ*. The log likelihood of any direction of motion can be computed from any other two non-degenerate log likelihoods, 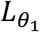 and 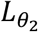:

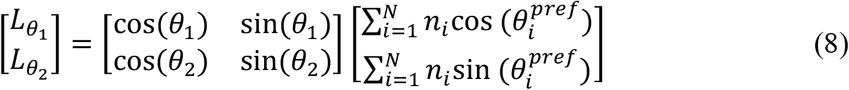

From this equation, the log likelihood of any other direction *θ*_3_ can be computed as follows:

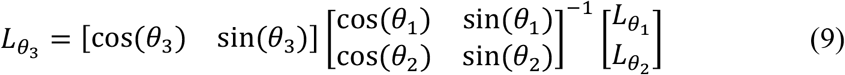

In each trial, we calculated the log likelihoods for 0° and 45° 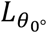 and 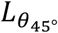, as 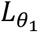 and 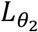 and then computed the log likelihoods of all other directions between –179° and 180° using 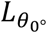 and 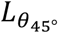. The direction with the largest log likelihood was used as an estimate of the stimulus motion, indicating the direction representation of the population response.

### Targeted dimensionality reduction (TDR)

To understand the effects of prior expectations on MT neural activity at the population level, we used targeted dimensionality reduction (TDR) method. A detailed explanation of TDR can be found in Mante et al. ^42^ For the population analysis, we constructed the population activity by pooling the responses of all the MT neurons across conditions (stimulus contrast, prior block, and target direction) in each monkey (Pursuit task: 137 neurons from Monkey A and 120 neurons from Monkey B; Fixation task: 82/86 neurons in the high/low contrast cases from Monkey A and 83/80 neurons in the high/low contrast cases from Monkey B). In the pursuit data, although the prior directions and the outer directions (± 15° and ± 120° from the prior direction) were not the same across sessions, the relative relationships between the prior direction and outer directions were the same. Therefore, a pseudo population was composed by aligning each neuron’s responses relative to the prior direction and combining them, meaning that each neruon’s preferred direction was redefined depending on the difference between the preferred direction and the prior direction.

During the pursuit task, targets in prior directions were presented in both the wide and narrow prior blocks but other targets in the outer directions were presented either in wide or narrow prior blocks. Therefore, we used linear regression with the neural responses to only the prior direction to describe the population responses as a linear combination of task variables (stimulus contrast and prior):

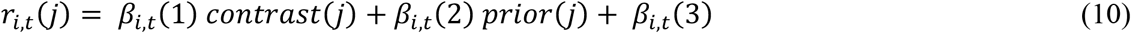

where *r*_*i,t*_ *(j)* is the z-scored response of neuron *i* at time *t* on trial j (mean and standard deviation are computed from the neuron’s responses across all trials and times); *contrast(j)* is the stimulus type on trial *j* (1: high contrast; 0: low contrast); *prior(j)* is the block type on trial *j* (1: wide prior block; 0: narrow prior block); *β*_*i,t*_(1) and *β*_*i,t*_ (2) are the regression coefficients that reflect the extent to which the trial-by-trial firing rate of neuron *i* at time *t* dependents on the corresponding task variable, contrast and prior, respectively; *β*_*i,t*_ (3) is the regression coefficient responsible for the trial-by-trial variation in firing rates due to time rather than task variables.

On the other hand, during the fixation task, all stimulus motions in 12 directions (0°, 30°, …, 300°, 330°) were presented in both prior blocks across all sessions. Therefore, the MT neural responses to all motion directions were used to compute the regression coefficients for contrast, prior and stimulus motion:

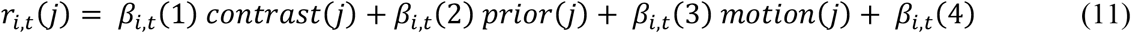

where *r*_*i*,t_*(j)* is the z-scored response of neuron *i* at time *t* on trial j; *contrast(j)* is the stimulus type on trial *j*; *prior(j)* is the block type on trial *j*; *motion(j)* is the motion direction of the stimulus (angles in radians); *β*_*i*,t_(1∼3) are the regression coefficients of neuron *i* at time *t* for stimulus contrast, prior, and motion direction of the stimulus, respectively; *β*_*i*,t_(4) is the regression coefficient of neuron *i* at time *t*, which captures the differences in firing rates across time.

We denoised the regression coefficients by projecting them into the population subspace spanned by the first 12 principal components from PCA, and then represented the denoised coefficients on the original basis (Figs. 5d, e, Supplementary Figs. 3d, e and Supplementary Figs. 4a-f). Next, we estimated the fixed regression vectors by time-averaging and orthogonalizing the regression coefficients. The fixed regression vectors were defined as task-related subspace axes that independently accounted for the trial-by-trial variance of population response due to task variables (contrast and prior for the pursuit task; contrast, prior, and stimulus motion for the fixation task). The average population responses were projected onto these axes to investigate the temporal dynamics of the population responses in the task-related neural subspace.

Additionally, we performed the TDR analysis on the pursuit task data using all motion direction conditions (the prior and outer directions) such that stimulus motion was included as a task variable. The analytical method was identical to that described above and the conclusions were similar (Supplementary Figs. 4g-t).

## Acknowledgments

We thank Ui hwan Moon and SeungHwan Lee for help with animal care, and Heeyeon Ahn for help with data preprocessing. We thank Dr. Stephen G. Lisberger and Hayoung Song for their helpful discussions and comments on the manuscript. This research was supported by the IBS-R015-D1. The authors declare no competing financial interests.

---

**Supplementary Fig. 1:**
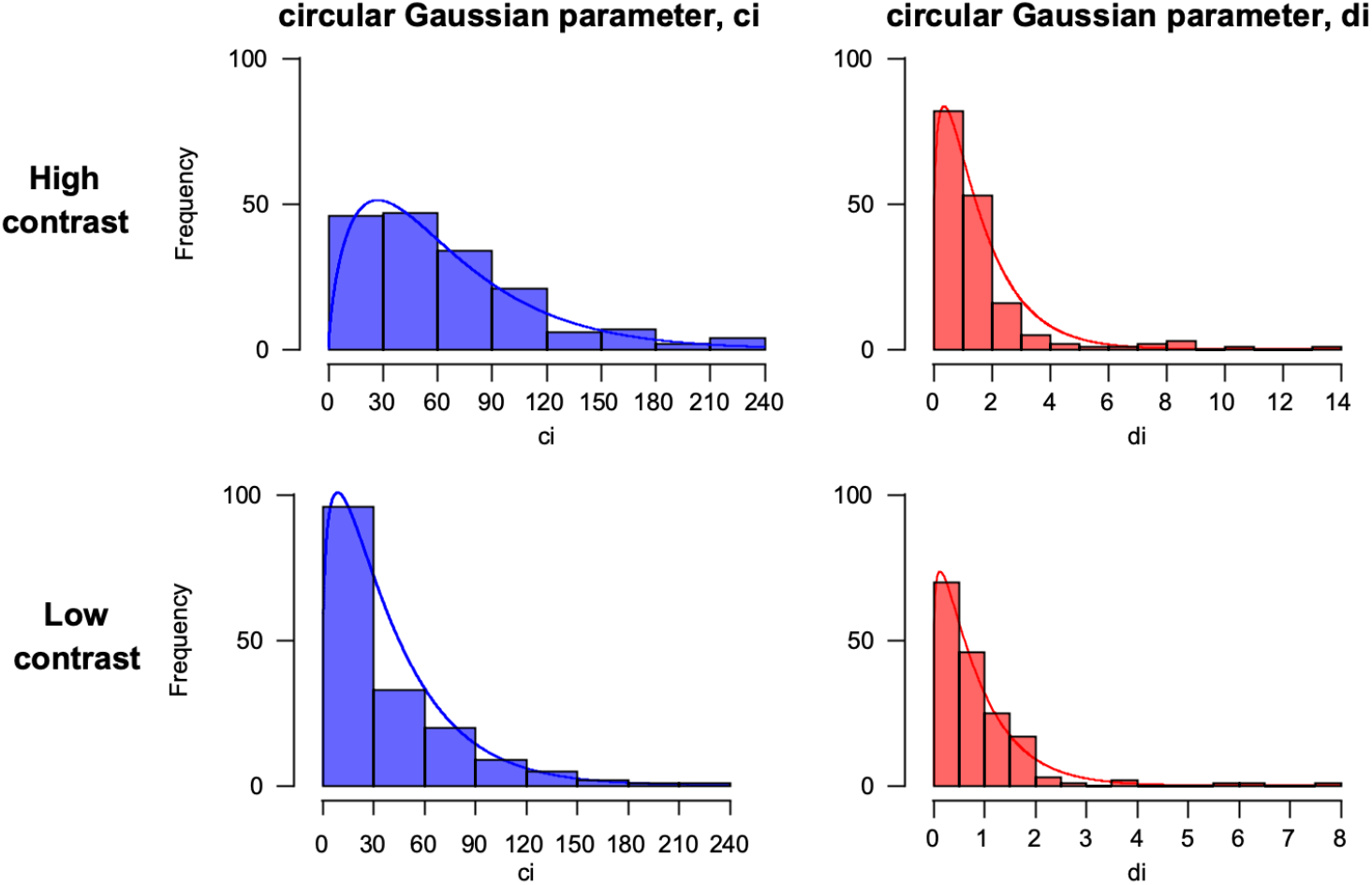
Distributions of circular Gaussian parameters. Distributions of *c*_*i*_ (blue) and *d*_*i*_ (red), which are the parameters of the circular Gaussian function fitted to the responses of *i*th recorded MT neuron: 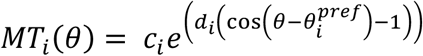, where *θ* is the 12 motion directions (0°, 30°, …, 330°) of the visual stimulus, and 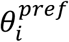 is the mean of the circular Gaussian function. The histograms show the frequency of the parameters estimated from the recorded data. The lines indicate the gamma distribution fitted to the estimated circular Gaussian parameters. The parameters (*c*_*i*_′ and *d*_*i*_′) are randomly selected from the gamma distribution for *i*th simulated MT neuron: *c*′_*i*_ ∼ Γ *(k*_*c*_, *θ*_*c*_*), d*′_*i*_ ∼ Γ *(k*_*d*_, *θ*_*d*_), where *k*_*c*_ and *θ*_*c*_ are the shape and scale parameters of the gamma distribution of *c*; *k*_*d*_ and *θ*_*d*_ are the shape and scale parameters of the gamma distribution of *d*, respectively.

**Supplementary Fig. 2:**
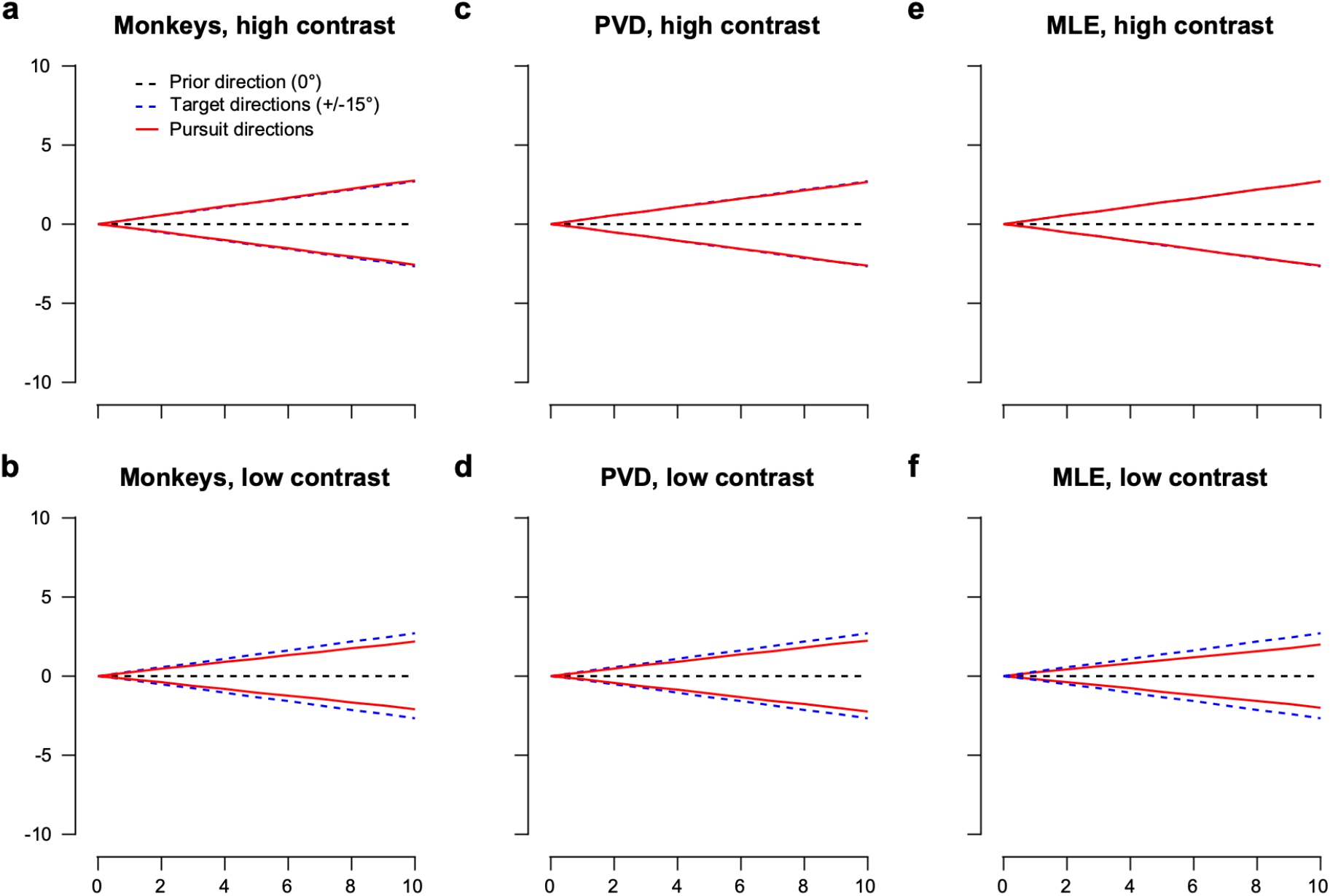
Experimental and simulated pursuit directions for outer target directions. The black dotted line indicate the prior direction and the blue dotted lines indicate the outer directions. The red solid lines indicate the average pursuit directions for the two outer directions in high (a, c, e) and low (d, e, f) contrast cases of the narrow prior block. The ratio between the angular difference in the two outer target directions and that in the corresponding pursuit directions quantifies the effect of prior expectation on the bias of pursuit direction traces ^35^. **a, b** Average pursuit directions from the combined dataset of the two monkeys. All prior directions were rotated to 0°, thus, the outer directions were aligned to ±15°. Only the days with more than 50 trials for each condition were included (46 days for Monkey A and 43 days for Monkey B) **c– f** Average pursuit directions estimated from simulated MT neuronal responses to ±15° directions using PVD (c, d) or MLE (e, f). Both experimental and simulated pursuit directions are biased in the prior direction (0°) by a similar amount.

**Supplementary Fig. 3:**
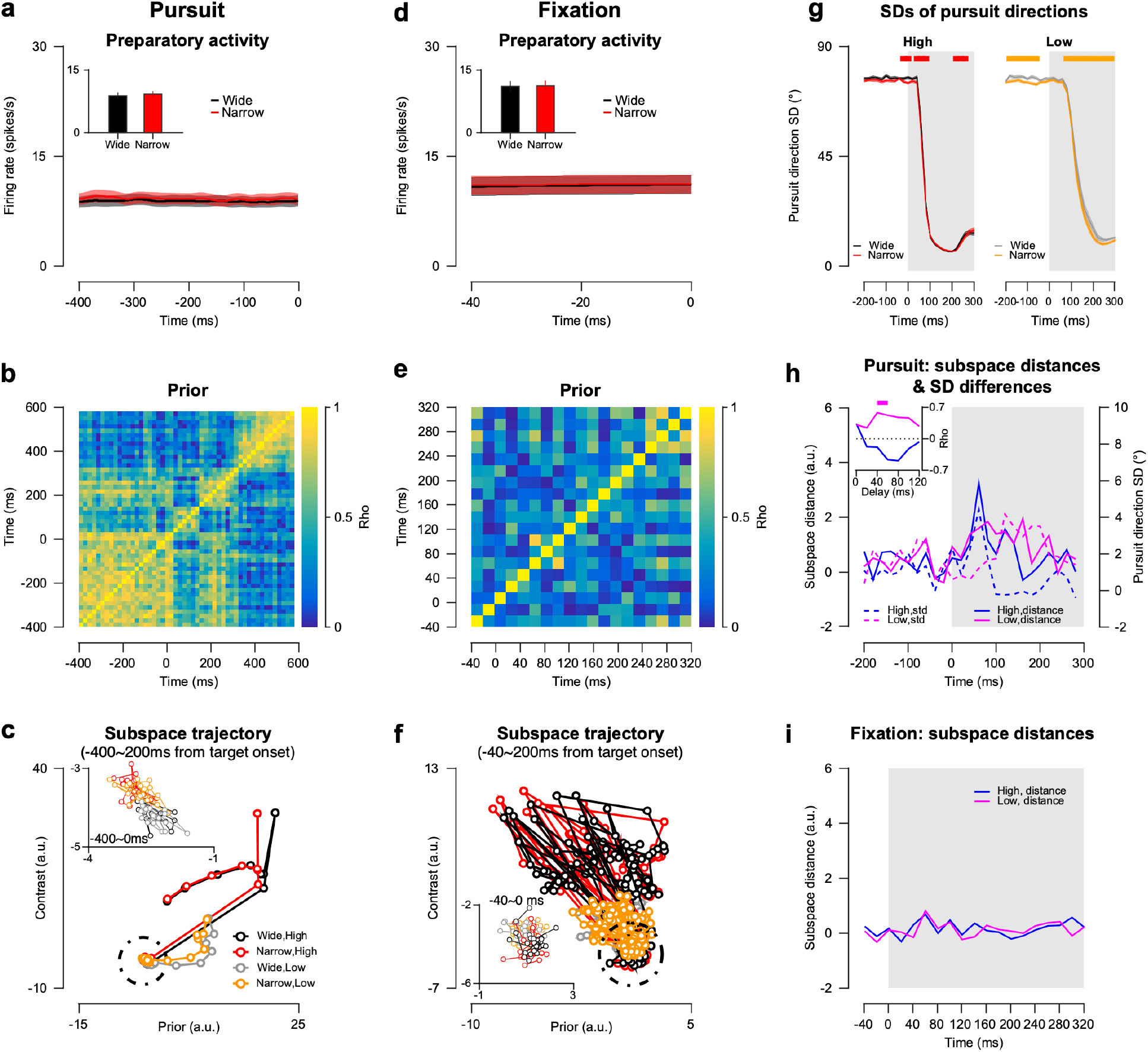
Temporal dynamics of MT population responses in Monkey B during pursuit and fixation task. Same as Fig. 5 but for Monkey B. **a-c** for the pursuit task. **a** PSTHs and mean preparatory responses of the MT neurons (inset). **b** Autocorrelation matrix of regression coefficients for prior. **c** Temporal trajectories of average population responses in the task-related subspace using the prior and contrast as axes. **d-f** Same as a-c but for the fixation task. **g** SDs of pursuit directions with high (left) and low (right) contrast stimuli as a function of time relative to the target onset. **h** Distances of prior-axis projected population responses between the wide and narrow prior blocks as a function of time (solid lines), and the differences in pursuit direction SDs between the two priors as a function of time (dotted lines). The blue and magenta lines indicate the high contrast and low contrast conditions, respectively. Temporal coherence between subspace distances and SD differences with different latencies of SD differences from subspace distances is shown in the inset plots. The thick top line in the inset indicates that the FDR-corrected p-values are less than 0.05. **i** Distances of population responses between the two prior blocks on the prior axis of the subspace during the fixation task.

**Supplementary Fig. 4:**
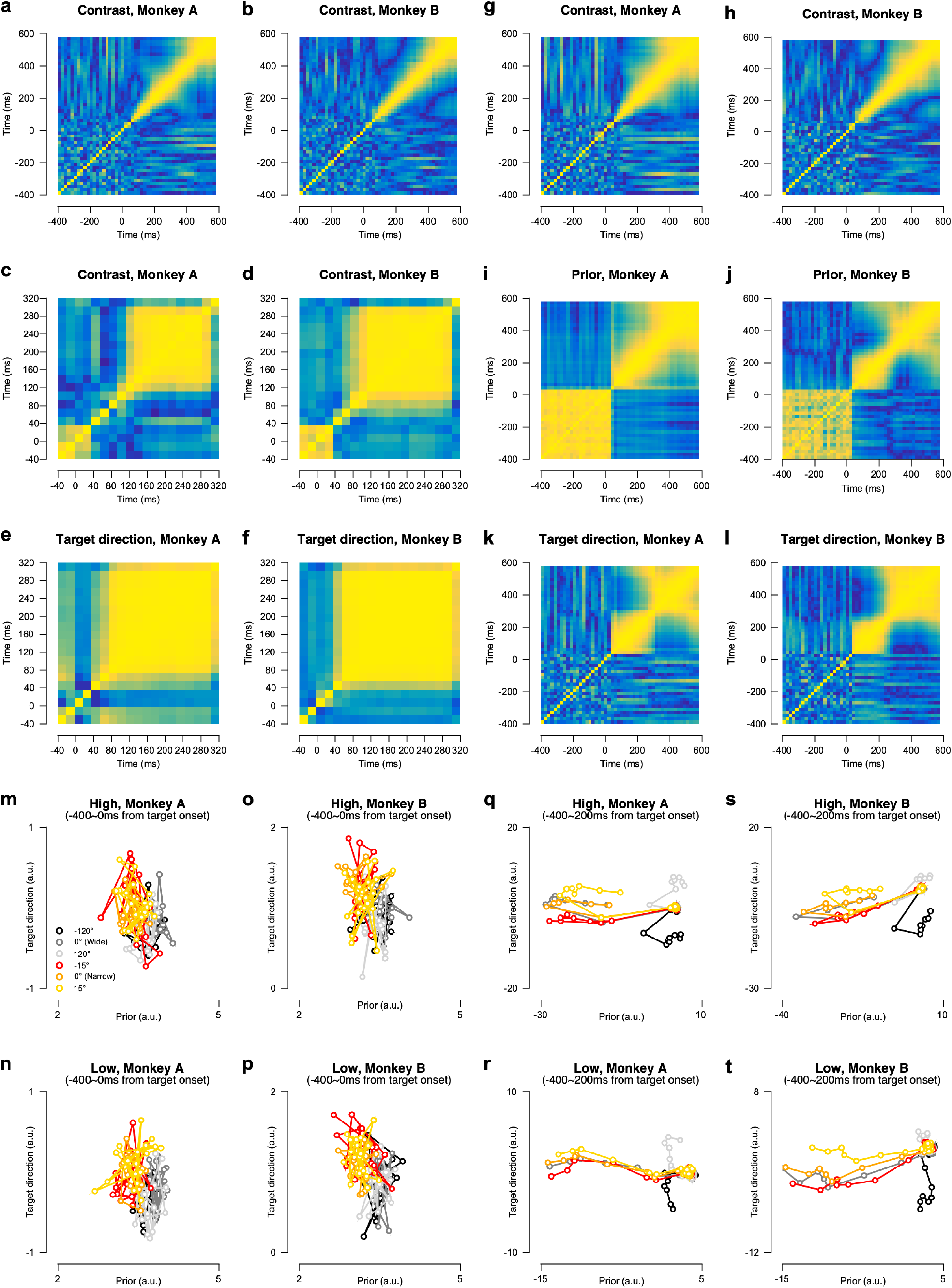
Temporal dynamics of MT population responses for other properties. **a, b** Autocorrelation matrix of regression coefficients for contrast during the pursuit task in Monkey A and B, respectively. **c, d** Autocorrelation matrix of regression coefficients for contrast during the fixation task in Monkey A and B, respectively. Temporal generalization of regression coefficients for contrast before stimulus onset is due to the stimulus presentation sequence in the fixation task. Four or five of stimuli with random directions were presented in one trial, but the stimulus contrast was identical in each trial. Therefore, monkeys were aware of the contrast in the stimulus sequence in each trial, which contributed to the generalized pattern of spontaneous activity. **e, f** Autocorrelation matrix of regression coefficients for target direction during the fixation task in Monkey A and B, respectively. **g-l** Autocorrelation matrix of regression coefficients for contrast (**g, h**), prior (**i, j**), target direction (**k, l**) during the pursuit task, where the regression coefficients were calculated from the population responses to all the target directions (–120°,–30°,0°,30°,120°). **m-t** Temporal trajectories of average population responses in task-related subspace using prior and target direction as axes. Trajectories of preparatory population responses in the subspace (**m-p**) and trajectories of population responses in the subspace before and after pursuit target onset (**q-t**).

**Supplementary Fig. 5:**
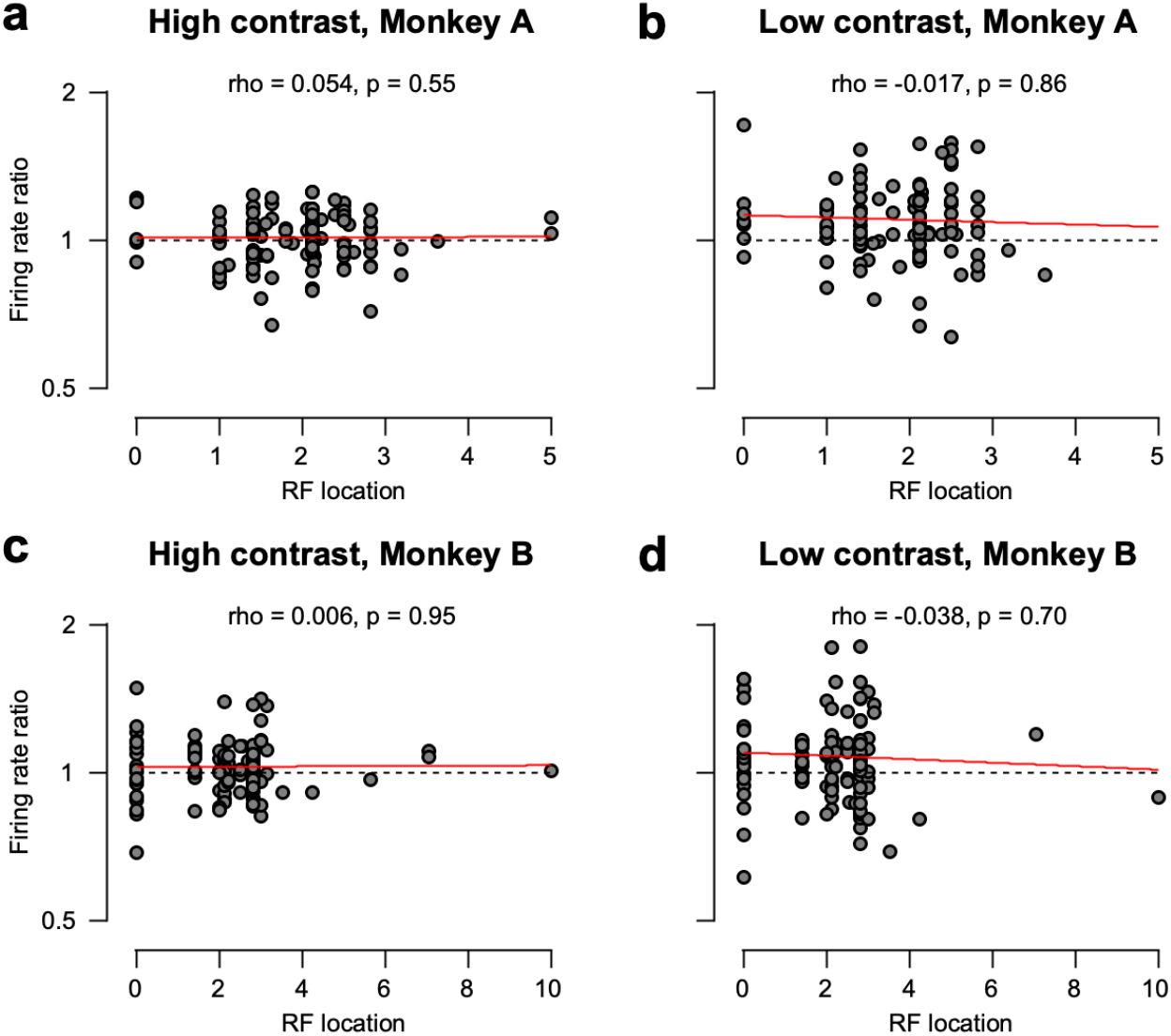
Relationship between the response reduction and receptive field location. **a, c** Relationship between the distance of each neuron’s receptive field from the central fixation point (RF location) and the ratio of firing rate in the narrow prior block to that in the wide prior block (firing rate ratio) for the high contrast case. The data points in the plot represent the firing rate ratio of MT neurons between the two prior blocks at each RF location. The red line represents the linear regression line of the RF location on the firing rate ratio. **b, d** Relationship between the receptive field location and firing rate ratio for the low contrast case: no significant relationship between the two under all the conditions for either monkey.

**Supplementary Fig. 6:**
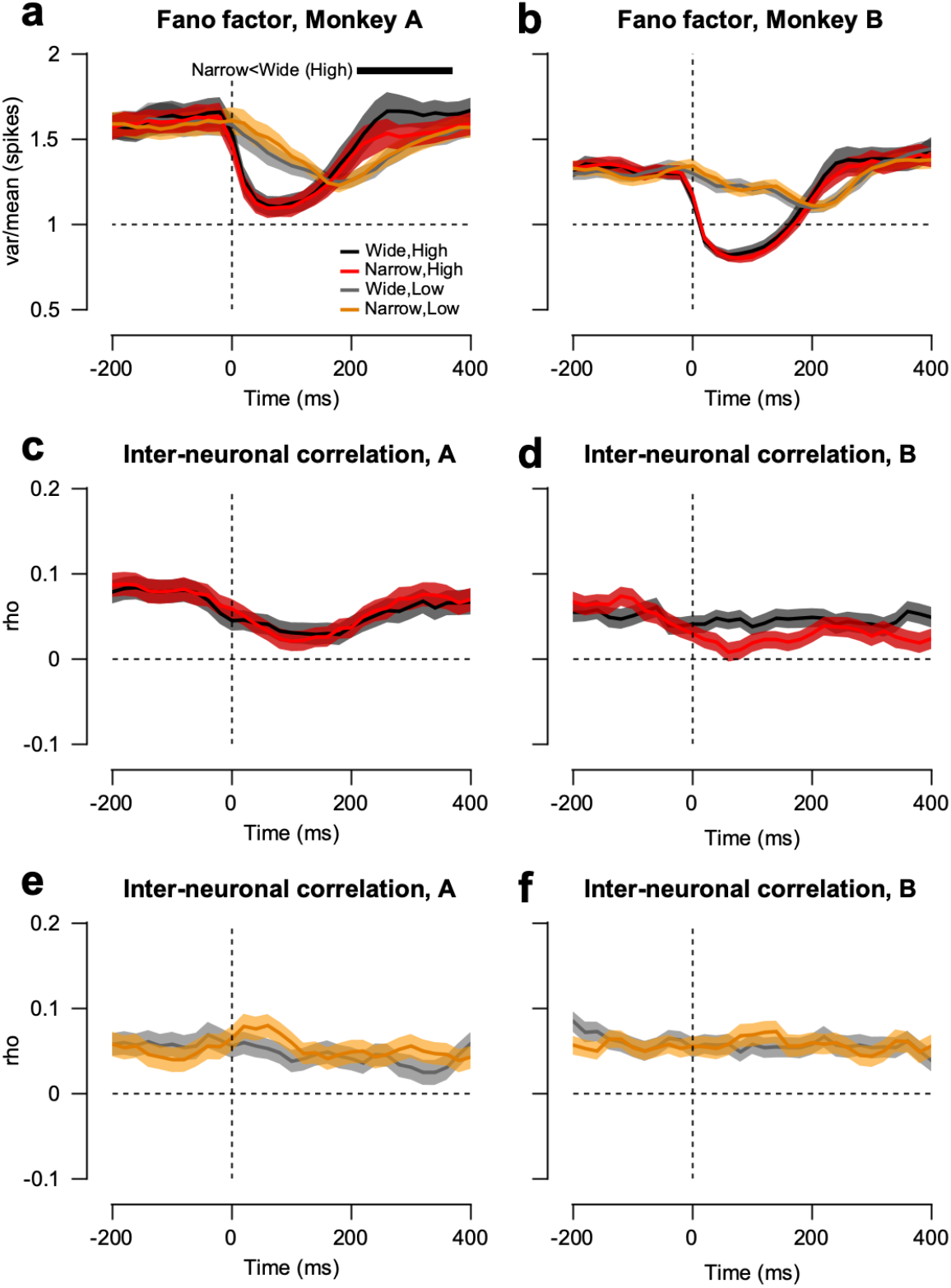
Spike count variability and inter-neuronal correlation. To calculate the spike count variability and inter-neuronal correlation as a function of time relative to the pursuit target onset, 100 ms (±50 ms) sliding time windows were used from –200 to 400 ms after the stimulus onset with a step size of 20 ms. **a, b** Mean normalized variance (Fano factor) of spike counts over time under four different conditions. The black and red lines show the mean Fano factors for high contrast cases in the wide and narrow prior blocks, respectively. The gray and yellow lines show the mean Fano factors for the low contrast cases in the wide and narrow prior blocks, respectively. Colored error bands indicate the standard error. **c, d** Pairwise spike count correlation as a function of time in the high contrast condition. **e, f** Pairwise spike count correlation as a function of time in the low contrast condition.

